# Multiomics Integration Elucidates Metabolic Modulators of Drug Resistance in Lymphoma

**DOI:** 10.1101/2021.01.07.425721

**Authors:** Fouad Choueiry, Satishkumar Singh, Xiaowei Sun, Shiqi Zhang, Anuvrat Sircar, Hart Amber, Lapo Alinari, Epperla Narendranath, Robert Baiocchi, Jiangjiang Zhu, Lalit Sehgal

## Abstract

**Background:** Diffuse large B-cell lymphoma (DLBCL) is the most common non-Hodgkin lymphoma (NHL). B-cell NHLs rely on Bruton’s tyrosine kinase (BTK) mediated B-cell receptor signaling for survival and disease progression. However, they are often resistant to BTK inhibitors or soon acquire resistance after drug exposure resulting in the drug tolerant form. The drug tolerant clones proliferate faster, have increased metabolic activity, and shift to oxidative phosphorylation; however, how this metabolic programming occurs in the drug resistant tumor is poorly understood.

**Methods:** In this study, we explored for the first time the metabolic regulators of ibrutinib-resistant activated B-cell (ABC) DLBCL using a ‘multi-omics’ analysis that integrated metabolomics (using high-resolution mass spectrometry) and transcriptomic (gene expression analysis). Overlay of the unbiased statistical analyses, genetic perturbation and pharmaceutical inhibition, were further used to identify the key players that contribute to the metabolic reprograming of the drug resistant clone.

**Results:** Gene-metabolite integration revealed interleukin 4 induced 1 (IL4I1) at the crosstalk of two significantly altered metabolic pathways involved in the production of various amino acids. We showed for the first time that drug resistant clones undergo metabolic reprogramming from glycolysis towards oxidative phosphorylation & is activated via the BTK-PI3K-AKT-IL4I1 axis and can be targeted therapeutically.

**Conclusions:** Our report shows how these cells become dependent on PI3K/AKT signaling for survival after acquiring ibrutinib resistance and shift to sustained Oxidative phosphorylation, additionally we outline the compensatory, pathway that regulates this metabolic reprogramming in the drug resistant cells. These findings from our unbiased analyses highlight the role of metabolic reprogramming during drug resistance development. Furthermore, our work demonstrates that a multi-omics approach can be a powerful and unbiased strategy to uncover genes and pathways that drive metabolic dysregulation in cancer cells.

## Introduction

Diffuse large B-cell lymphoma (DLBCL) is the most common non-Hodgkin lymphoma (NHL). The majority of patients with DLBCL (~60%) are cured with Rituximab-Cyclophosphamide, Hydroxydaunomycin, Oncovin and Prednisone (R-CHOP) chemotherapy, but nearly 1 in 3 patients will eventually relapse.[1, 2] Patients with relapsed/refractory disease have a high rate of morbidity and mortality despite treatment with salvage therapies. Patients with DLBCL can be classified into either the activated B-cell (ABC) molecular subtype or the germinal center B-cell (GCB) molecular subtype based on gene expression profiling and are characterized by distinct gene mutation signatures.[3] Following multi-agent chemotherapy, the ABC molecular subtype has a lower survival rate compared to the GCB subtype.[4, 5] Because ABC-DLBCL has chronically active B-cell receptor signaling, several components of these signaling pathways are attractive therapeutic targets.[3] Bruton’s tyrosine kinase (BTK) drives the B-cell receptor-signaling cascade, which leads to the activation of NF-kB and other targets.[6, 7] The orally administered, bioavailable BTK inhibitor ibrutinib has been FDA-approved to treat patients with relapsed mantle cell lymphoma (MCL), Waldenstrom’s macroglobulinemia, and chronic lymphocytic leukemia (CLL), including those harboring 17p deletion.[8, 9] In a phase I/II clinical trial of relapsed/refractory DLBCL, 37% of ABC-DLBCL patients responded to ibrutinib.[10] Despite these encouraging results, ibrutinib treatment produces variable or incomplete responses and leads to drug-resistance and genetic alterations stemming from unknown causes.[10, 11]

Therapeutic targeting metabolic alterations has emerged as a promising strategy for hematological malignancies.[12] Molecular subsets of lymphomas possessing distinct metabolic signatures provide the unique opportunity to investigate the role of metabolic changes in disease progression and response to chemotherapeutics. These lymphomas are categorized based on cellular reliance on increased mitochondrial oxidative phosphorylation versus cellular reliance on B-cell receptor signaling.[13] Previously, Pera et al. (2018) described the metabolic consequences of a lysine deacetylase inhibitor (KDACI) in DLBCL and revealed KDACI-induced lymphoma cell dependency on choline metabolism. This multi-omics approach has also been used to study the metabolic dysregulations impeding necroptosis in DLBCL[14], paving the way for the identification of metabolic vulnerabilities in cancer.[12]

Metabolomics can identify metabolic changes or anomalies in biological systems and deliver qualitative and semi-quantitative information about the steady-state prevalence of substrates from various metabolic pathways. Thus, metabolomics studies can provide an overview of the metabolic dysregulations affecting cellular physiology in disease. The metabolome reflects the phenotypic outcome of genomic, transcriptomic, and proteomic influences on the molecular state of the cell. For this reason, the metabolome can be considered the end readout of ‘multi-omics’-driven cell physiology. In the context of cancer biology, the activation of oncogenes by genetic mutation or deregulated gene expression can drive a cancer cell into a physiological state in which cell metabolism is effectively reprogrammed. A genomics approach can identify genetic mutations that can derail the cell’s physiology, while a transcriptomic approach can identify RNA species whose abnormal expression contributes to an altered physiological state of the cancer cell. When multiple ‘omics’ profiles are interrogated in parallel, the resulting data can be integrated into a more complete “story” about the cells under study. In summary, a multi-omics approach provides a multi-layered insight into how upstream changes (such as those observable in genomic, transcriptomic, and proteomic profiles) exert changes on downstream metabolic indicators of the physiological state of the cell.

Here, we performed an untargeted metabolomics workflow using a QExactive high-resolution mass spectrometer to investigate metabolic reprogramming associated with ibrutinib resistance in paired ibrutinib -sensitive and ibrutinib -resistant DLBCL cell lines.[15] We also performed a transcriptome analysis of these cell lines to identify genes expressed at different levels in the ibrutinib -sensitive and ibrutinib -resistant cells. Multi-omics analysis of the metabolome and transcriptome data identified common denominators, indicative of candidate molecular pathways driving ibrutinib resistance in DLBCL.

## Results

### Establishing Ibrutinib-resistant DLBCL lines

Ibrutinib-resistant clones of the DLBCL cell lines HBL1 and TMD8 were generated in vitro as reported previously[15] and named HBL1R and TMD8R hereafter. Activated B-cell lymphoma pathogenesis exhibits irregular initiation of the BTK–mediated B-cell receptor signaling pathway[16] (Figure 1A). Ibrutinib, a selective and irreversible BTK inhibitor, has been successfully utilized in various leukemia and lymphoma models.[17] The culture media was perpetually supplemented with ibrutinib to induce a resistant phenotype of the cells (Figure 1B). Only cells with fewer than 20 passages were used for experiments. Wild type cells, along with their newly induced resistant clones, were challenged with Ibrutinib to confirm a shift in drug tolerance (Figure 1C and Figure 1D). Half-maximal inhibitory concentration (IC50) assays of the lymphomas cultured in incremental doses of ibrutinib confirm the acquired drug-resistance in these DLBCL lymphomas. HBL1R had a minimum inhibitory concentration of 11733nM, while the sensitive cells were only tolerant at 851.5nM. Similarly, TMD8R had an increased inhibitory concentration of Ibrutinib (9661nM) compared to the parental sensitive TMD8 cells (9.226nM). Ibrutinib resistance appeared to be accompanied by a more rapid proliferation rate in the HBL1R and TMD8R cell lines compared to their ibrutinib -sensitive parental lines. To validate this observation, the growth rate of different DLBCL cells was measured (Figure 1E and Figure 1F). The steeper slopes for the resistant cells in the growth curves reflect their faster proliferation rates compared to the sensitive cells. A significant difference in growth rates became evident at 48 hours for the HBL1/HBL1R pair (p-value= 0.013). The difference in TMD8/TMD8R growth was quickly apparent at 24 hours after seeding (p-value= 0.00072). Various studies have revealed that the adaptation of metabolic processes is essential to sustained proliferation in cancer cells.[18] Considering this, our observations of increased proliferation and decreased doubling time prompted the hypothesis that altered metabolic processes in ibrutinib - resistant DLBCL cells are switched on to promote cellular growth and sustained or increased resistance to Ibrutinib. In the following sections, we interrogated the HBL1/HBL1R paired cell lines for metabolomic and transcriptomic differences. We subsequently tested a subset of identified markers in the TMD8/TMD8R paired cell lines as a means of validation.

**Figure 1:**
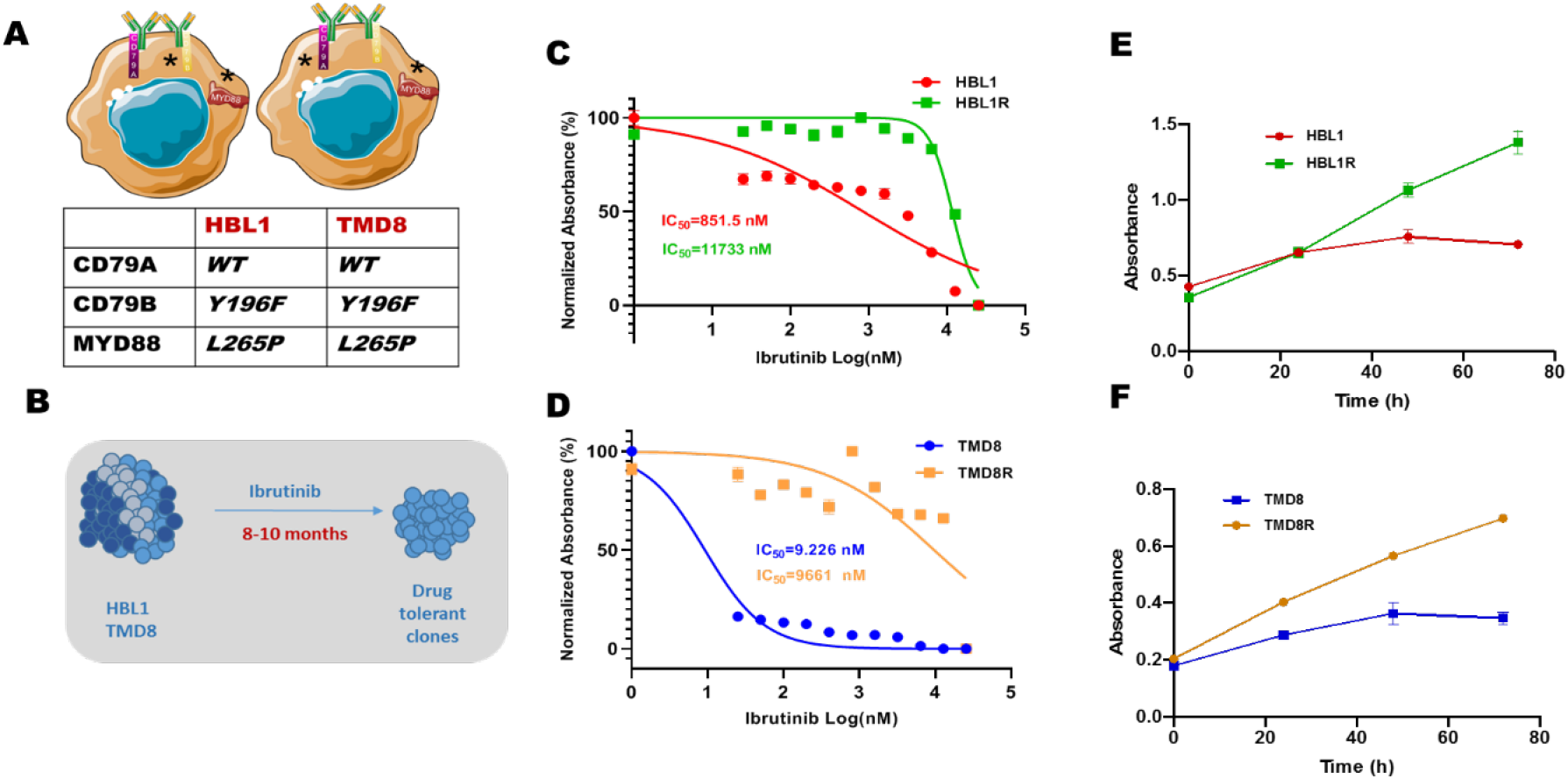
Generation of the ABC Ibrutinib-resistant DLBCL cell lines. A) Wild-type HBL1 and TMD8 cells were selected for their B cell receptor (BCR) independence and MD88 gene expression (Y196F – L265P?). B) HBL-1 and TMD8 cells were perpetually cultured in Ibrutinib to induce drug resistance. C-D) Half-maximal inhibitory concentration (IC50) graphs of the lymphomas cultured in incremental doses of Ibrutinib. E-F) Proliferation assay to determine cell growth in HBL1R and TMD8R compared to their respective wild-type parental lines.

### Determining the Metabolic Modifications Accompanying the Ibrutinib Resistance Phenotype of HBL1

To characterize the metabolic profile of ibrutinib - resistant cells, we performed a comprehensive untargeted metabolomics analysis with extensive compound identification processes using multiple databases including the human metabolite database (HMDB), Kyoto encyclopedia of genes and genomes (KEGG) and PubChem compound database, and our in-house high-resolution mass spectra database. Our analysis identified 603 intracellular polar metabolites mutually detected in both HBL1 and its resistant clone. Partial least square discriminant analysis (PLS-DA) was performed to observe the metabolic differences between the two cell phenotypes. As demonstrated, the biological replicates of the two cell lines are tightly clustered within their groups ensuring good biological reproducibility. More importantly, the experimental groups are separated from each other validating their distinct metabolic profile (Figure 2A). PLS-DA component 1 (x-axis) highly summarized 47.6% of the variance, while 13.9% of variations (y-axis) of our detected metabolites are explained by component 2. The top 15 metabolites driving separation of the clusters in the PLS-DA plot were identified and showed in the variance importance in the projection (VIP) plot (Figure 2B). A greater VIP score indicates the higher importance of these metabolites in driving separation of the drug-resistant and drug-sensitive cell lines.

**Figure 2:**
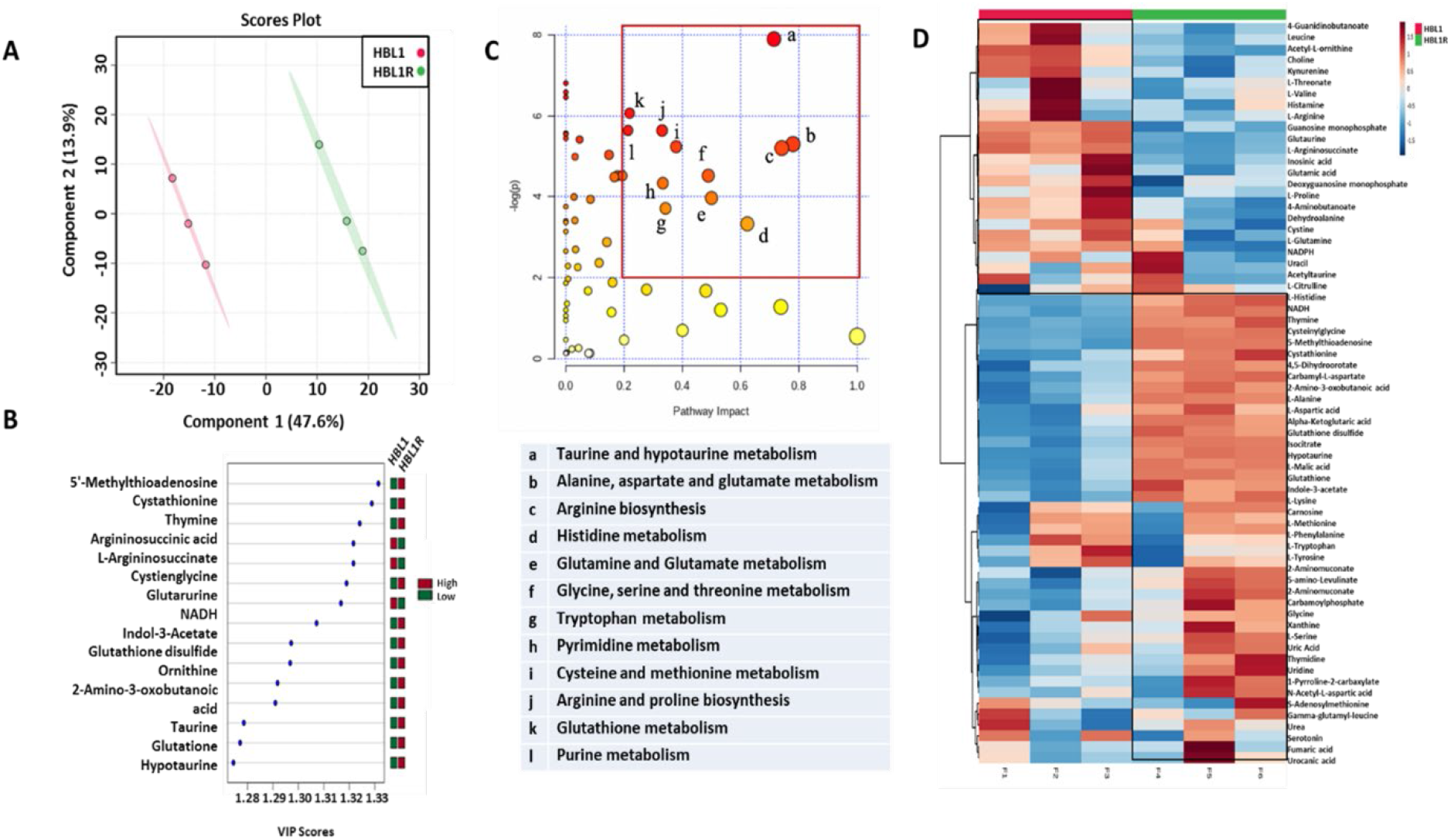
Metabolic profiling of HBL1. A) PLS-DA plot depicting the variability in metabolic profiles of HBL1 versus its resistant clone, HBL1R. B) VIP plot showing the metabolites driving separation of the uniqueness in metabolic profiles. C) Pathways analysis revealing the top dysregulated metabolic pathways across the two phenotypes. D) Heatmap of metabolites from significantly altered pathways revealing changes in levels among groups.

The color map on the right of the VIP plot of Figure 2B shows the relative quantity of these metabolites. The top three metabolites driving the separation of the drug-resistant and drug-sensitive clusters were identified as 5’-methylthioadenosine, cystathionine, and thymine. As these individual metabolites partake in complex metabolic processes, a summary of the major altered pathways was generated (Table 1). The twelve most impacted metabolic pathways were selected based on the impact score > 0.2 and −log(p) > 2 (Figure 2C). The taurine and hypotaurine metabolic pathways were most greatly affected, and purine and pyrimidine metabolism were revealed to be among the most dysregulated pathways. However, amino acid metabolic pathways dominated the list. Metabolites from the top altered pathways (e.g., arginine metabolism, proline metabolism, and pyrimidine metabolism) were subsequently selected for further statistical analysis.

**Table 1:** Significantly altered metabolic pathways in HBL1R phenotype.

| Pathway Name | p | FDR | Impact |
| --- | --- | --- | --- |
| Alanine, aspartate and glutamate metabolism | 4.98E-03 | 2.40E-02 | 7.80E-01 |
| Arginine and proline metabolism | 3.56E-03 | 2.40E-02 | 3.30E-01 |
| Arginine biosynthesis | 5.50E-03 | 2.40E-02 | 7.41E-01 |
| Cysteine and methionine metabolism | 5.31E-03 | 2.40E-02 | 3.78E-01 |
| Glutamine and glutamate metabolism | 1.89E-02 | 5.18E-02 | 5.00E-01 |
| Glutathione metabolism | 2.32E-03 | 2.40E-02 | 2.19E-01 |
| Glycine, serine and threonine metabolism | 1.09E-02 | 3.61E-02 | 4.89E-01 |
| Histidine metabolism | 3.57E-02 | 7.27E-02 | 6.23E-01 |
| Purine metabolism | 3.55E-03 | 2.40E-02 | 2.14E-01 |
| Pyrimidine metabolism | 1.31E-02 | 3.99E-02 | 3.33E-01 |
| Taurine metabolism | 3.72E-04 | 2.27E-02 | 7.14E-01 |
| Tryptophan metabolism | 2.44E-02 | 5.95E-02 | 3.42E-01 |

At the crosstalk of various pathways reported in Figure 2C, sixty-eight metabolites from the significantly dysregulated pathways in HBL1R cells were detected and the results were compared to HBL1 cells. Average metabolite abundance from both HBL1 and HBL1R cells was log-transferred and their fold change ratio is described in Supplemental Table 1. The relative abundance of the sixty-eight metabolites is shown as a heatmap in Figure 2D. In this figure, each column depicts a biological replicate from the experiment while rows convey the relative abundance of individual metabolites. The red color shows relatively high expression while blue indicates lower expression. The heatmap demonstrates that the differences in individual metabolite levels from the two types of cells (HBL1 and HBL1R) are strikingly significant. Statistical analysis identified twenty-nine significantly altered metabolites among the sixty-eight metabolites (p-value < 0.05). Of those twenty-nine, fifteen metabolites including prominent pathway intermediates from cysteine and methionine metabolism and urea cycle (e.g., cystathionine and argininosuccinate, respectively) significantly drive separation of the groups (p-value < 0.001). Additionally, the major tricarboxylic acid (TCA) cofactor NADH was detected among the top metabolites involved in the significantly altered metabolism (p-value = 0.0005).

### Discovering a Genetic Network Driving Altered Metabolic Pathways using Multiomics Integration

While untargeted metabolomics provided a plethora of data, it is representative of a biological snapshot of the metabolic profiles of the cells at the time of harvest.[19] To further enable mechanistic understanding, transcriptomic data from the HBL1 and HBL1R cell lines were also generated to further validate the observed metabolic changes. As reported in Figure 3A, gene set enrichment analysis (GSEA) of DNA microarray data was conducted to observe changes of genes along various metabolic pathways based on the curated KEGG gene set from the molecular signatures database. A total of 169 dysregulated gene sets between the HBL1 and HBL1R were found, of which 99 were upregulated in the resistant cells. A false discovery rate (FDR) cutoff of 0.3 was used to increase confidence in the data, rendering 58 significant gene sets (Table 2) that may be significantly altered when HBL1 cells acquire drug resistance.

**Figure 3:**
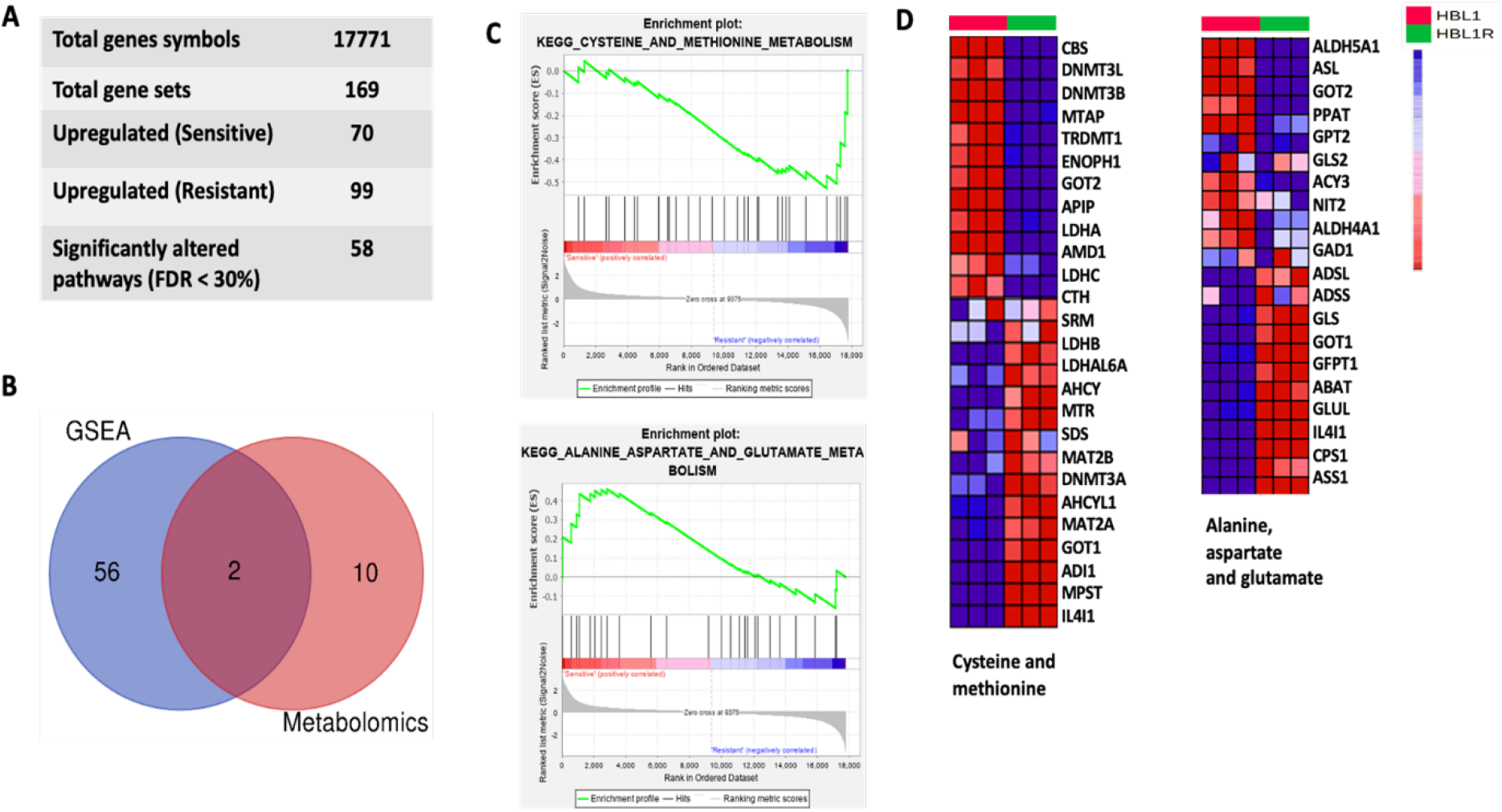
Integration of altered pathways for HBL1 sensitive and resistant clones. A) Summary of the gene set enrichment analysis of HBL1/HBL1R expression data from the DNA microarray. B) Overlap of gene and metabolite data shows two altered metabolic pathways at both the metabolic and transcriptional levels. C) GSEA plot of HBL1/R highlighting the overlapping dysregulated pathways in both HBL1 datasets. D) Heatmap indicating the altered genes in cysteine and methionine as well as alanine aspartate and glutamate metabolic pathways.

**Table 2:**
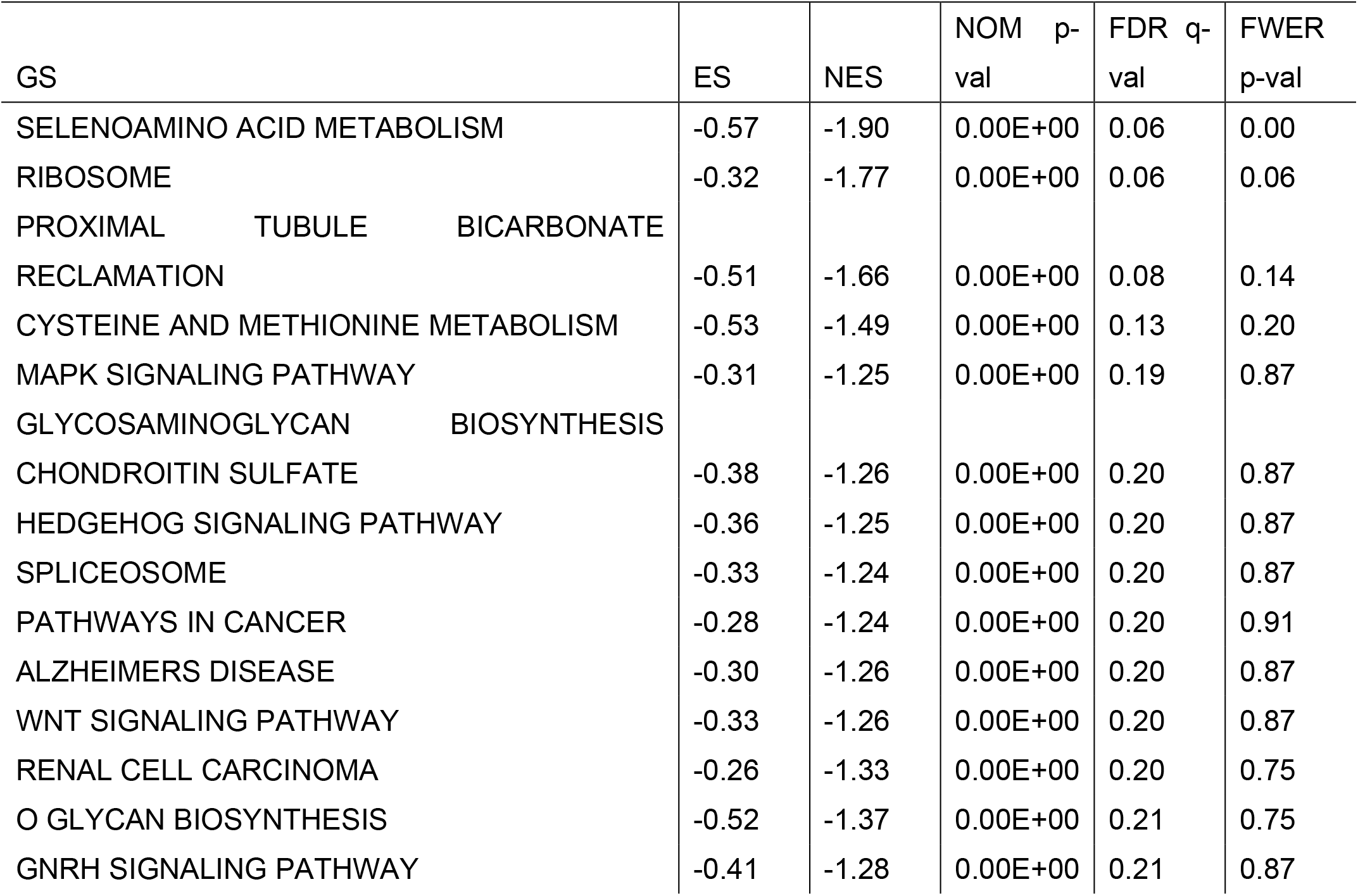
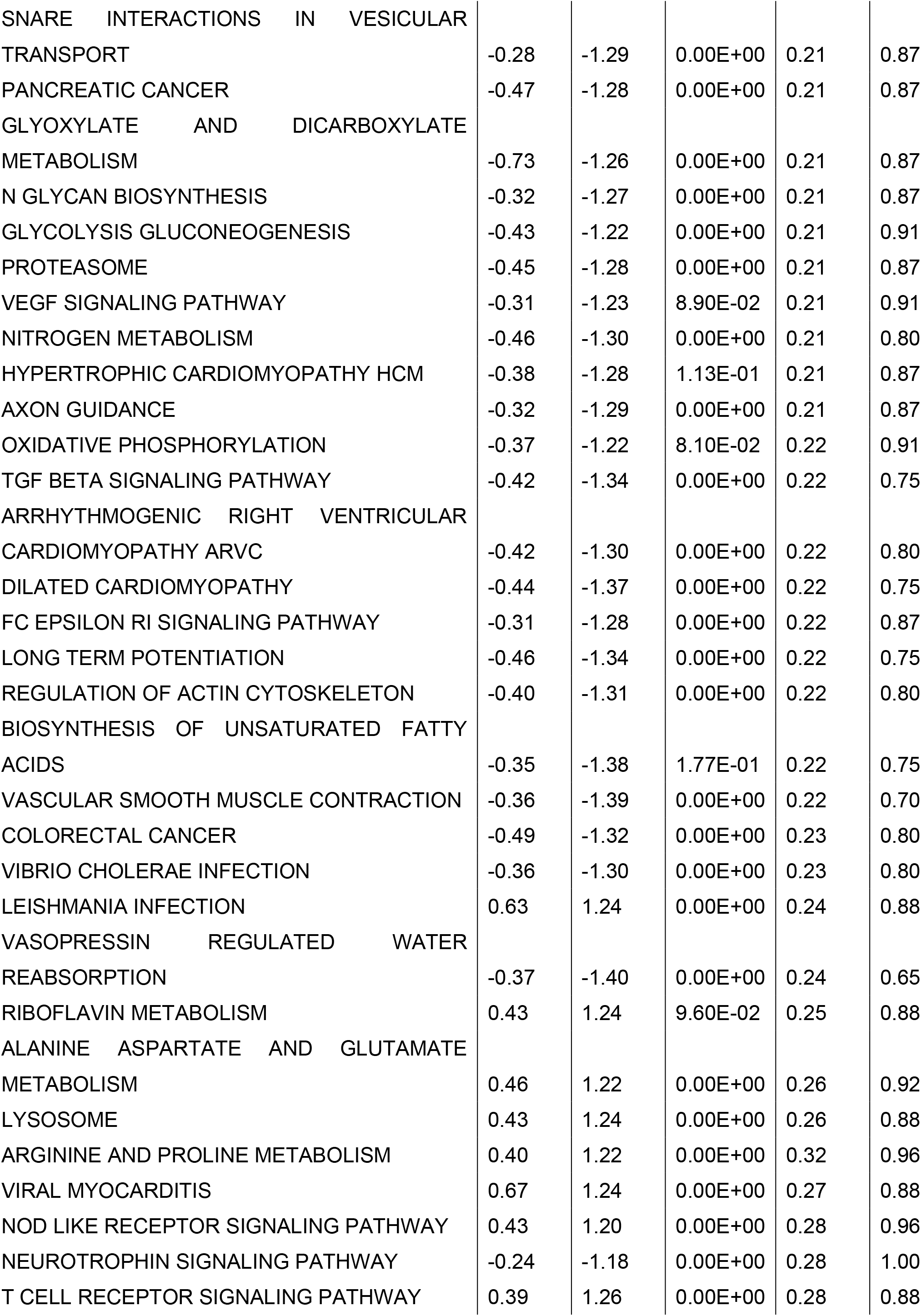
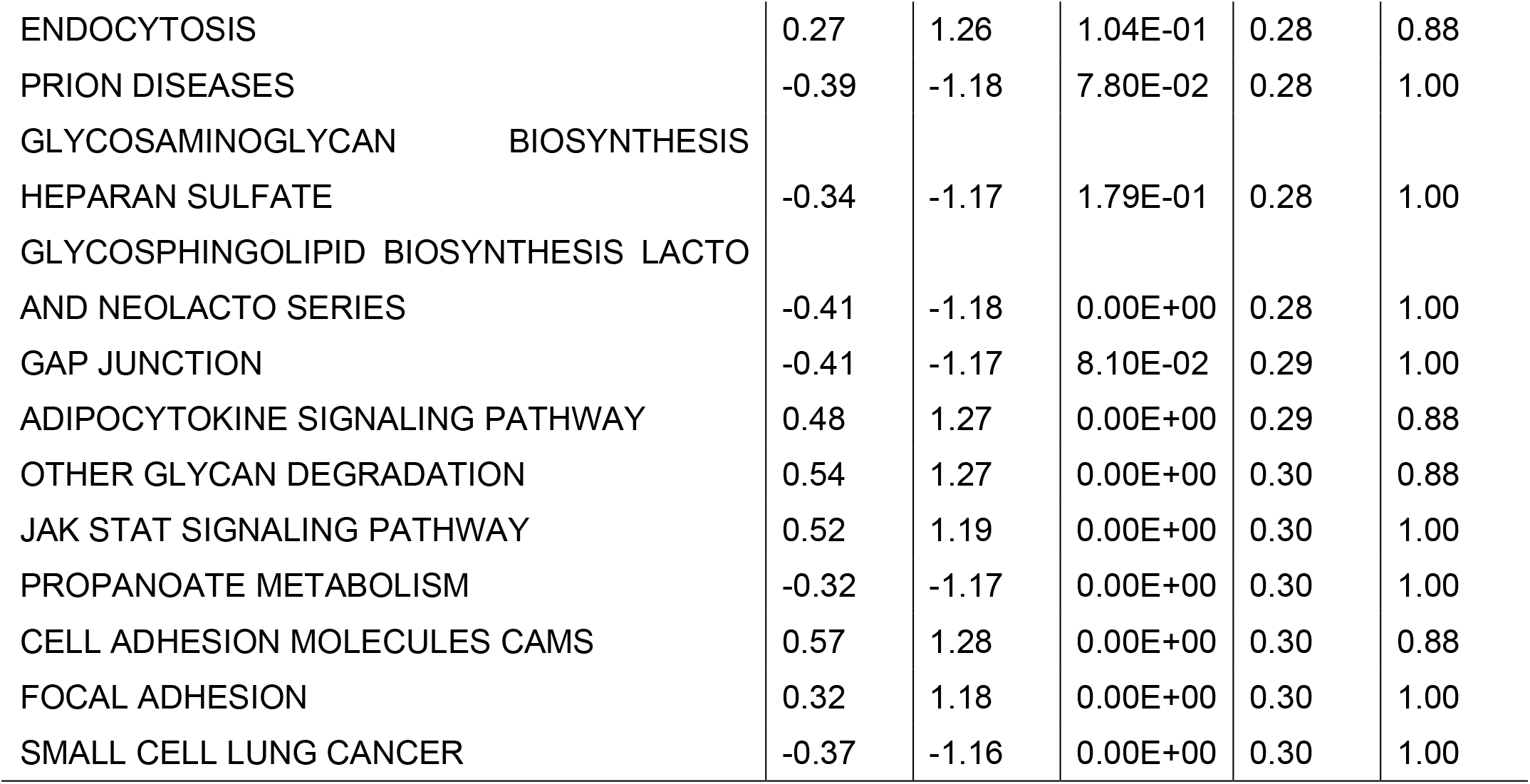
Significantly altered microarray KEGG gene sets in HBL1R phenotype.

Further investigation was performed to determine the significant alterations observed at both the metabolic and transcriptional levels. The overlap of the top fifty-eight altered pathways from microarray GSEA along with the twelve dysregulated pathways from untargeted metabolomics analysis highlighted that there were two metabolic pathways significantly dysregulated at both the transcript and metabolite level (Figure 3B). The two detected pathways encompass (1) cysteine and methionine metabolism, and (2) alanine, aspartate, and glutamate metabolism. GSEA analyses revealed alanine, aspartate, and glutamate metabolism to be down-regulated in the resistant phenotype; conversely, cysteine, and methionine metabolism was found to be upregulated (Figure 3C). The heatmap presentation of gene fold changes detected from these pathways is depicted in Figure 3D. Two genes that are involved in cystathionine synthase and lyase (CBS and CTH, respectively) and that belong to the cysteine and methionine metabolism KEGG pathway are expressed at higher levels in HBL1 than in HBL1R. The genes IL4I1 and MPST were the most downregulated in HBL1R compared to HBL1. For the alanine, aspartate, and glutamate metabolism KEGG pathway, argininosuccinate 1 (ASS1) and carbamoyl phosphate 1 (CPS1) were the most downregulated genes in HBL1 compared to HBL1R.

### Multi-omics Integration Highlights Metabolic Shift Towards Oxidative Phosphorylation

To further investigate these metabolic changes and to confirm the observed trend in Ibrutinib-resistant DLBCL metabolism, we identified genes overlapping in both dysregulated pathways and quantified their expression via q-PCR. Altered genes of each unique pathway were identified and reported in Supplemental Table 2 and 3. Twenty-six genes were detected from the cysteine and methionine metabolic pathways (Supplemental Table 2), and twenty-four genes were detected from alanine, aspartate, and glutamate metabolic pathways (Supplemental Table 3). The enzymes Interleukin 4 induced 1 (IL4I1) and glutamic-oxaloacetic transaminase 1 and 2 (GOT1 and GOT2) were implicated in both sets of metabolic pathways (Figure 4A). IL4I1 belongs to the L-amino-acid oxidase (LAAO) family and catalyzes the oxidation of L-phenylalanine to keto-phenylpyruvate.[20] The IL4I1 protein also works in conjunction with additional amino-transferases to target other amino acids. As shown in Figure 4B, IL4I1 mRNA expression was analyzed by q-PCR and found at significantly decreased levels in HBL1R (p-value= 0.014) and TMD8R (p-value= 0.0052) compared to wild-type HBL1 and TMD8, respectively. GOT1 and GOT2 mRNA expression levels were significantly decreased in HBL1R (p-value= 0.013) and TMD8R (p-value=0.02), respectively. Additionally, genes levels of drug resistance targets ALDH7A1, ALDH9A1, and LAP3 were diminished in both resistant cells. Gene-metabolite interactions from our two dysregulated pathways were analyzed and mapped (Figure 4C). This interaction network revealed that IL4I1 links methionine and aspartic acid, thereby bridging the gap between cysteine and methionine metabolism and alanine, aspartate, and glutamate metabolism. These proteins mediate the reversible transamination from glutamate to oxaloacetate, generating α-ketoglutarate and aspartate.[21] In turn, those metabolites form a direct interaction with 4-aminobutyrate aminotransferase (ABAT), linking alanine, α-ketoglutarate, and gamma-aminobutyric acid (GABA) into this drug-resistance mechanism. Lymphomas are generally considered glycolytic.[22] As reprogramming of energy metabolism is a hallmark of cancer progression[23], we questioned whether a shift to oxidative phosphorylation (OXPhos) occurred in the drug-resistant clone. Kaplan-Meier survival plots were generated for DLBCL patients using the publicly accessible R2 genomics visualization platform.[24] Gene Expression Omnibus data repository number GSE31312 was used to analyze IL4I1 levels to determine if there is a relationship between disease outcome and gene expression.[25, 26] Decreased expression of IL4I1 was directly correlated with lower chances of survival (p-value=0.019). IL4I1 expression can thus be linked with a more aggressive type of lymphoma.

**Figure 4:**
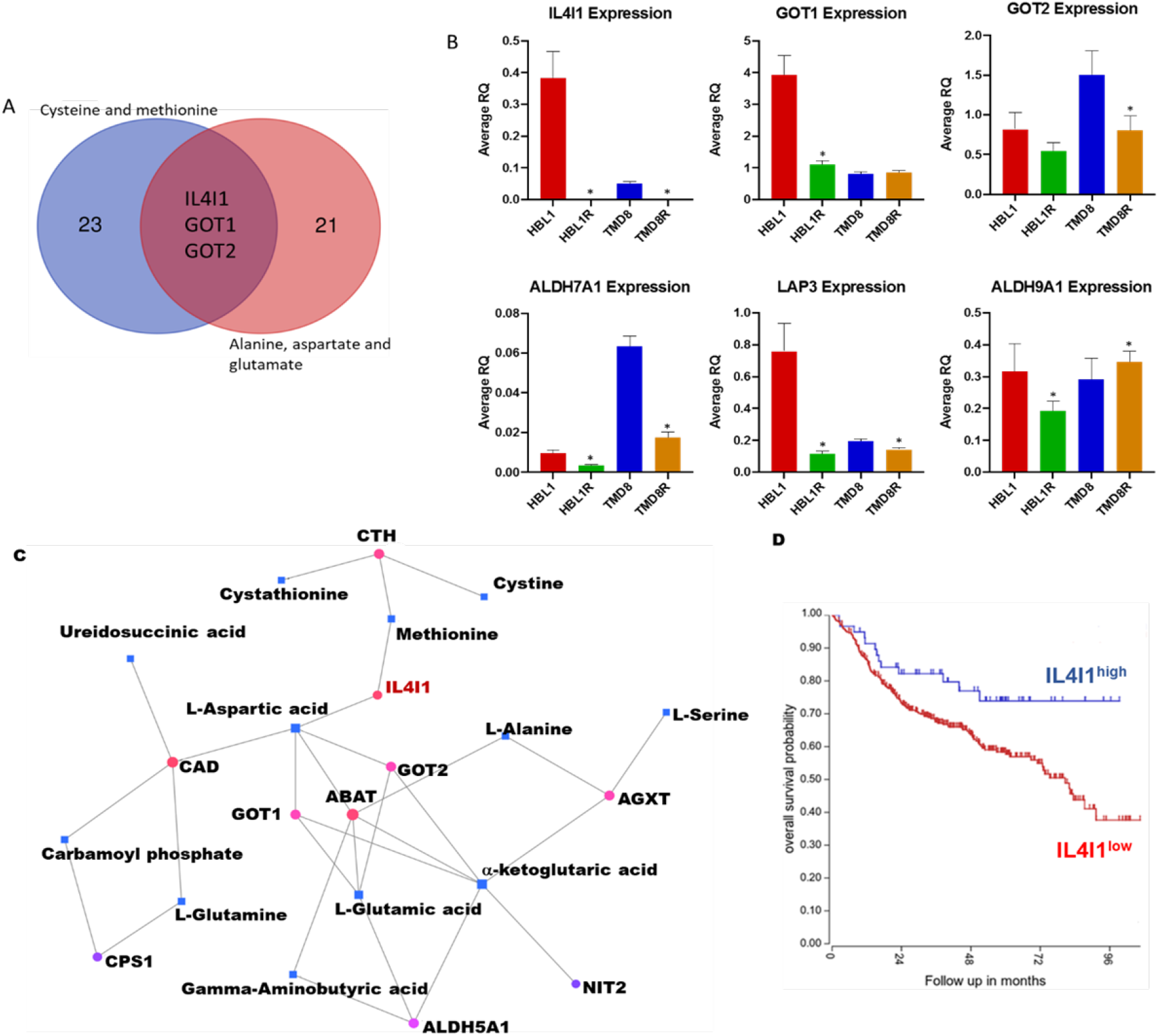
Integration of metabolic genes highlight IL4I1. A) Overlap of the genes pertaining to each metabolic pathway revealed IL4I1, GOT1 and GOT2 to be at the crosstalk of these pathways. B) q-PCR quantification of the genes of interest in lymphomas to validate expression. C) Proposed interaction map of transcriptomic and metabolomic data, hypothesized to be regulated by IL4I1. D) Kaplan-Meier survival analysis of DLBCL patients as determined from GSE31312 and analyzed by R2: Genomics Analysis and Visualization Platform.

### Metabolic Profiling of TMD8R Validates the Drug Resistance Related Metabolic Shift

To validate that the observed metabolic changes from HBL1 and HBL1R cells are truly indicative of a drug-resistant phenotype in the ABC subtype of DLBCL cells, untargeted metabolomics of a similarly induced Ibrutinib-resistant cell line of TMD8 (described here as TMD8R) was also conducted. In the TMD8 and TMD8R pairs of cells, 607 metabolites were detected. PLS-DA was performed to observe the separation of metabolic profiles from Ibrutinib-sensitive and resistant phenotypes (Figure 5A). From the PLS-DA score plot, component 1 was attributed to a 35.9% variation of all detected metabolites, while component 2 explains 28% of the variation. This conveys the distinct baseline metabolic profiles of each phenotype. Pathway analysis revealed fourteen significantly altered metabolic pathways in TMD8 and TMD8R pairs (Supplemental Figure 1A). The pathways were selected with identical inclusion criteria to that of the HBL1 and HBL1R pair, and are summarized in Table 3. Much like HBL1 and HBL1R, the TMB8/TMD8R pair also showed dysregulation of amino acid metabolic pathways. Three pathways were uniquely dysregulated in the TMD8/TMD8R pair: tyrosine metabolism, starch and sucrose metabolism, and glycerophospholipid metabolism. Changes in these three pathways may be attributed to the unique baseline metabolic profiles of this cell line. The dysregulated metabolic pathways in TMD8/TMD8R encompassed seventy-four metabolites (Supplemental Table 4). Variations in metabolite detection levels are represented in the heatmap in Supplemental Figure 1B. Twenty-eight of those pathway metabolites were significantly altered (p-value < 0.05), of which thirteen compounds (including fumarate, aspartate, and α-ketoglutarate) were significant with a p-value < 0.001.

**Figure 5:**
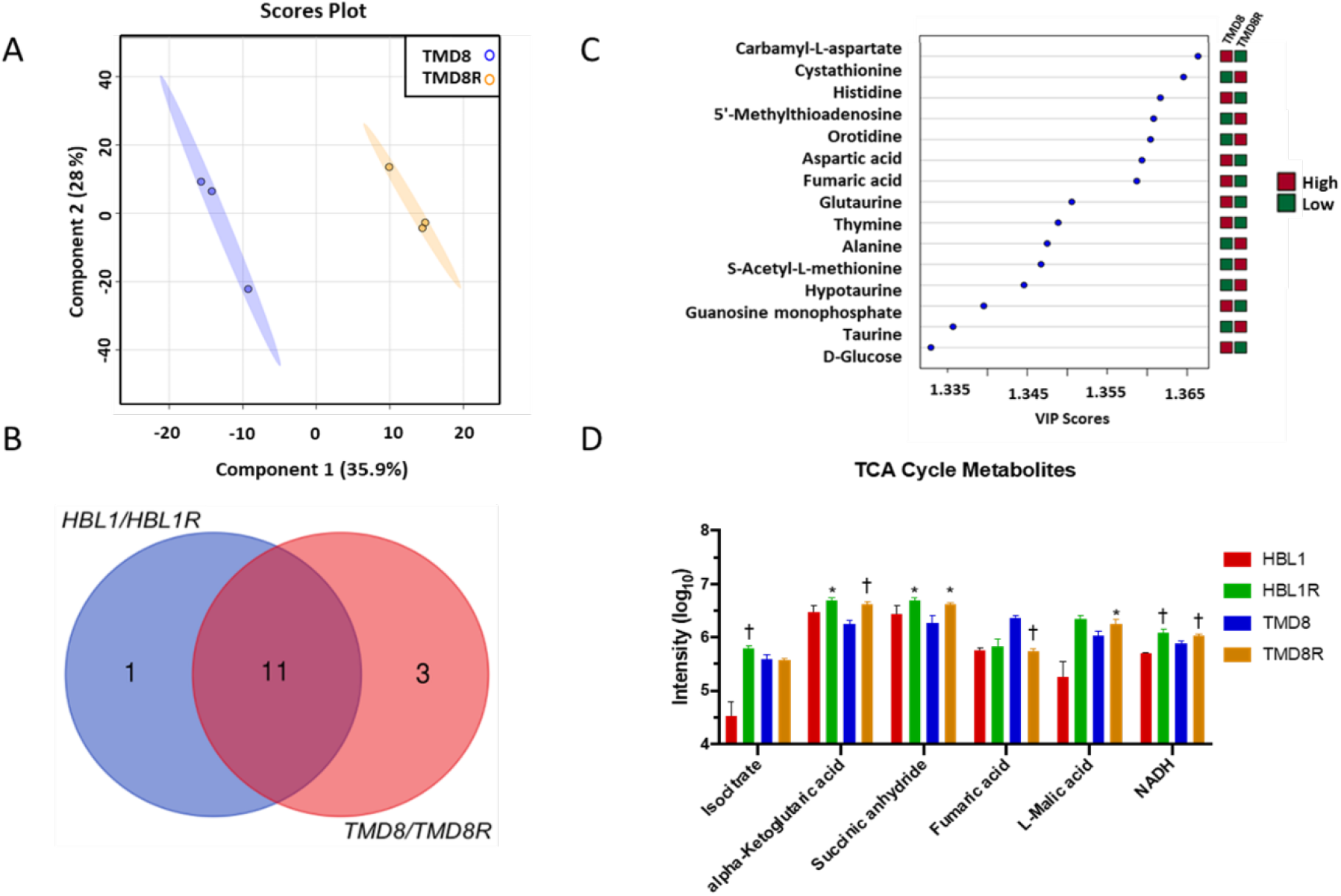
Validation of metabolic resemblance of TMD8R. A) PLS-DA plot depicting the variability in metabolic profiles of TMD8 versus its resistant clone, TMD8R. B) Overlap of top altered pathways in each cell line pairs post induced Ibrutinib resistance. C) VIP plot showing the metabolites driving separation of the uniqueness in metabolic profiles. D) Relative abundance of main altered metabolites for oxidative phosphorylation based on untargeted metabolomics analysis of each cell type.

**Table 3:** Significantly altered metabolic pathways in TMD8R phenotype.

| Pathway Name | p | FDR | Impact |
| --- | --- | --- | --- |
| Alanine, aspartate and glutamate metabolism | 3.85E-05 | 8.33E-04 | 7.80E-01 |
| Arginine and proline metabolism | 3.24E-02 | 6.92E-02 | 3.30E-01 |
| Arginine biosynthesis | 1.61E-02 | 4.76E-02 | 7.41E-01 |
| Cysteine and methionine metabolism | 2.70E-03 | 1.20E-02 | 3.78E-01 |
| D-Glutamine and D-glutamate metabolism | 2.49E-03 | 1.20E-02 | 5.00E-01 |
| Glutathione metabolism | 5.77E-03 | 2.03E-02 | 2.19E-01 |
| Glycerophospholipid metabolism | 6.37E-02 | 1.16E-01 | 2.77E-01 |
| Glycine, serine and threonine metabolism | 3.98E-04 | 3.53E-03 | 4.89E-01 |
| Histidine metabolism | 1.69E-04 | 2.63E-03 | 6.23E-01 |
| Purine metabolism | 5.28E-03 | 2.03E-02 | 2.14E-01 |
| Pyrimidine metabolism | 5.97E-04 | 4.11E-03 | 3.33E-01 |
| Starch and sucrose metabolism | 3.01E-02 | 6.66E-02 | 4.81E-01 |
| Taurine and hypotaurine metabolism | 2.45E-04 | 3.04E-03 | 7.14E-01 |
| Tyrosine metabolism | 8.07E-02 | 1.39E-01 | 4.00E-01 |

A comparison of metabolomic analyses between HLB1R and TMD8R revealed 11 dysregulated pathways that were shared by the two drug-resistant DLBCLs (Figure 5B). Much like the HBL1/HBL1R pair, metabolites from cysteine and methionine biosynthesis as well as alanine, aspartate, and glutamate metabolism were significantly dysregulated in the TMD8/TMD8R pair. Interestingly, alanine, aspartate, and glutamate metabolism was the top dysregulated pathway in TMD8/TMD8R. As purines and pyrimidines are the monomeric units of both DNA and RNA, the upregulation of their metabolism supports unrestrained development of tumors and may offer insight into increased proliferation.[27] The top fifteen metabolites with the highest VIP values that drive the separation of TMD8 and TMD8R cells in the PLS-DA plot were plotted in Figure 5C to identify similarities of metabolites pertaining to drug resistance. Six VIP metabolites were overlapping in both cell pairs, namely cystathionine, 5’-methylthioadenosine, glutaurine, thymine, hypotaurine, and taurine. The validation of Ibrutinib resistance-related metabolic similarities was achieved by the detection of overlapping metabolites driving the separation of Ibrutinib-sensitive and Ibrutinib-resistant phenotypes in both pairs of the lymphoma cell line.

To challenge the observation that Ibrutinib resistance in DLBCL cells shifts cellular energetics towards oxidative phosphorylation, we compared TCA metabolites (including the potent reducing agent NADH) of both pairs of drug-resistant and drug-sensitive cells (Figure 5D). Malic acid, α-ketoglutaric acid, and succinic anhydride were all detected at significantly higher concentrations in the resistant phenotypes (p-value < 0.05). An increase in isocitrate levels was also significantly detected (p-value < 0.001) but only in HBL1R. NADH, the prominent reducing agent involved in the TCA cycle, was detected at significantly increased levels in both resistant phenotypes (p-value < 0.001).

### Wild-type BTK expression restored IL4I1 in resistant tumors and Loss of IL4I1 contributes to resistance and metabolic reversal

The level of IL4I1 is downregulated in the ibrutinib resistant clones in both cell type at protein (Figure 6A) and mRNA level (Figure 4C). To further confirm if IL4I1 can contribute to the drug resistance, we knocked down IL4I1 in HBL1 and TMD8 parental cells at protein level using specific small interfering RNA (siRNA) targeting IL4I1, scrambled siRNA served as control (Figure 6B). Further, the loss of IL4I1 in the parental cells were able to elicit ibrutinib resistance similar to the acquired resistant clones (Figure 6C). To test whether the loss of BTK changes the levels of IL4I1 and affects Ibrutinib resistance, we ectopically expressed the wild-type (WT) BTK in Ibrutinib-resistant cells and measured expression of IL4I1. Compared to controls, IL4I1 levels increased upon ectopic expression of either WT-BTK or mutant C481S BTK (Figure 6D). Ectopic expression of wild-type BTK delayed cell proliferation (Figure 6E and 6F, Supplemental figure 2A-C). Wild-type BTK increased sensitivity of the HBL1R derived cells to Ibrutinib (Figure 6G) but not the TMD8R derived cells (Figure 6H). These results suggest that the expression of BTK can regulate the level of IL4I1 and contribute to Ibrutinib resistance. Our previous analysis showed that loss of BTK activates the PI3K-AKT pathway. To test whether IL4I1 expression is regulated by the PI3K-AKT axis, we treated the cells with BEZ235, a dual inhibitor of mTOR and AKT. As shown in Supplemental Figures 2B and 2C, the IL4I1 levels were restored after treating with the dual inhibitor, suggesting that IL4I1 is partly regulated by PI3K-AKT signaling. To further ascertain whether the restored IL4I1 levels in HBL1R-WTBTK affect the metabolism of the drug-resistant clone, untargeted metabolomics were performed on the clone. Our analysis revealed 458 intracellular polar metabolites mutually detected in the transduced mutants and annotated using the previously described spectral databases. To observe the metabolic similarities between the two cell phenotypes, PLS-DA was performed. PLS-DA plots revealed that the wild-type BTK expression group clustered adjacently to the wild-type HBL1 groups rather than HBL1R, suggesting a more similar metabolic profile to sensitive cells (Figure 6I). We found that the levels of key metabolites are reversed, such as α-ketoglutarate (p-value = 0.017), suggesting that the expression of BTK can reverse the expression of IL4I1 and the associated metabolic changes (Figure 6J-K). Similarly, this effect was also evident in TMD8-derived cells (Supplemental Figure 3A). TMD8R cells transduced with wild-type BTK (TRW) clustered further away from the parental TM8R cells and closer to the sensitive cells, suggesting a more similar metabolic pattern to TMD8. IACS-010759 selectively inhibits complex I of the mitochondrial electron transport chain (ETC), thereby disrupting OXPHOS, the metabolic process some tumor cells rely on for growth and survival[28]. Further, we found that the ibrutinib resistant cells HBL1R are now more sensitive to the treatment by IACS-010759 supplemental figure 3C suggesting that Metabolic reprogramming toward oxidative phosphorylation can be targeted therapeutically using OxPhos Inhibitors.

**Figure 6:**
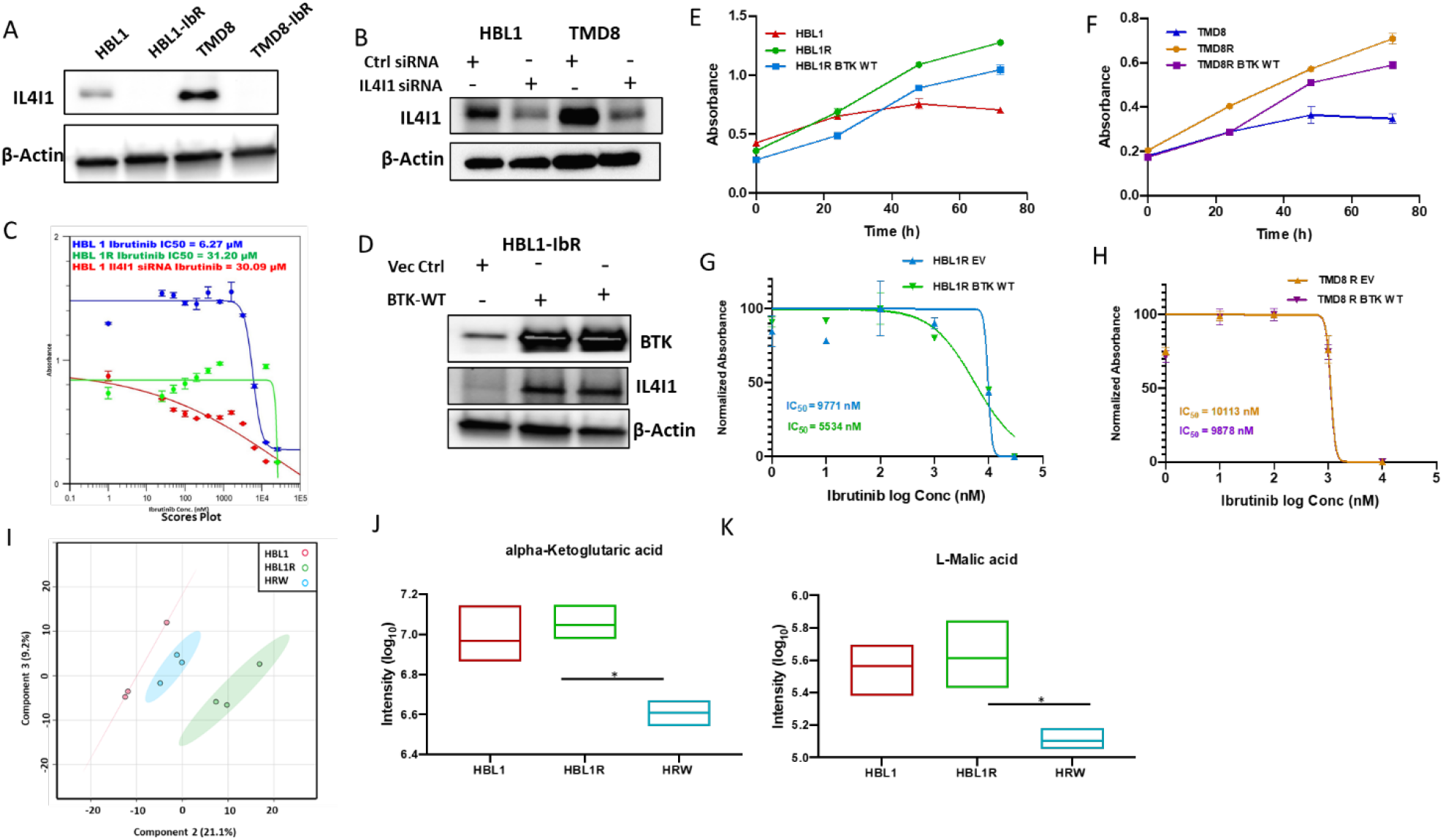
Wild-type BTK expression in the drug resistant cells hints at the partial metabolic reversal. A) Western blot of baseline IL4I1 expression across the lymphomas and their resistant clones. B) IL4I1 expression changes upon transfection by siRNA specific to IL4I1 or scrambled siRNA in HBL1 and TMD8 cells as shown by western blots. C) IC50 graphs of the HBL1 cells with either control (blue) or IL4I1 siRNA (red) and HBL1 resistant (green) lymphomas treated with Ibrutinib. HBL1 (IC50 6.2uM) HBL1 IL4I1 siRNA (IC50 30.08uM) HBL1R (IC50 31.20 uM). D) Western blot of IL4I1 expression upon ectopic expression of BTK WT in HBL1 cells. E-F) Proliferation assay to determine cell growth in the WT-BTK expressing resistant cells compared to their respective parental lines. G-H) IC50 graphs of the WT-BTK expressing resistant lymphomas treated with Ibrutinib. I) PLS-DA plot depicting the variability in metabolic profiles of HBL1, HBL1R, and HBL1R expressing WT-BTK. J-K) Relative signal intensity of TCA metabolites across the HBL1 derived cells.

### Proposed intersection of amino acid metabolism and TCA cycles within the Ibrutinib resistance mechanism regulated by IL4I1

As resistant lymphoma metabolism is established on an intersection of amino acid metabolic pathways, we continued our effort to establish a gene-metabolite network to help us identify a mechanism for the observed replenishing of the TCA cycle. While the mechanism by which IL4I1 can regulate metabolism remains unclear, we propose a data-driven network to explain this TCA anaplerosis. Potential routes linking these metabolic processes were constructed by mapping genes to metabolites through the biochemical pathways to which they pertain. As shown in Figure 7, the TCA cycle was centered with several key metabolites detected in our study, such as fumaric acid and α-ketoglutarate. These metabolites were focused on, as they open a door for potential direct metabolic links to take place. Highlighted on the right of Figure 7, IL4I1 is under the regulation of BTK. Our data suggest that decreased expression of IL4I1 will halt numerous amino acids from being converted to acetyl-CoA, the main product of glycolysis. This allows a link for amino acids, such as alanine, to be converted to pyruvate to sustain energy metabolism. Here, alanine aminotransferase 1 (GPT) and alanine-glyoxylate aminotransferase (AGXT) were both found to be upregulated (Supplemental Figure 4) in addition to increased abundance of alanine (p-value = 0.024). This provides a possible mechanism by which cells can propagate the TCA cycle in the absence of IL4I1.

**Figure 7:**
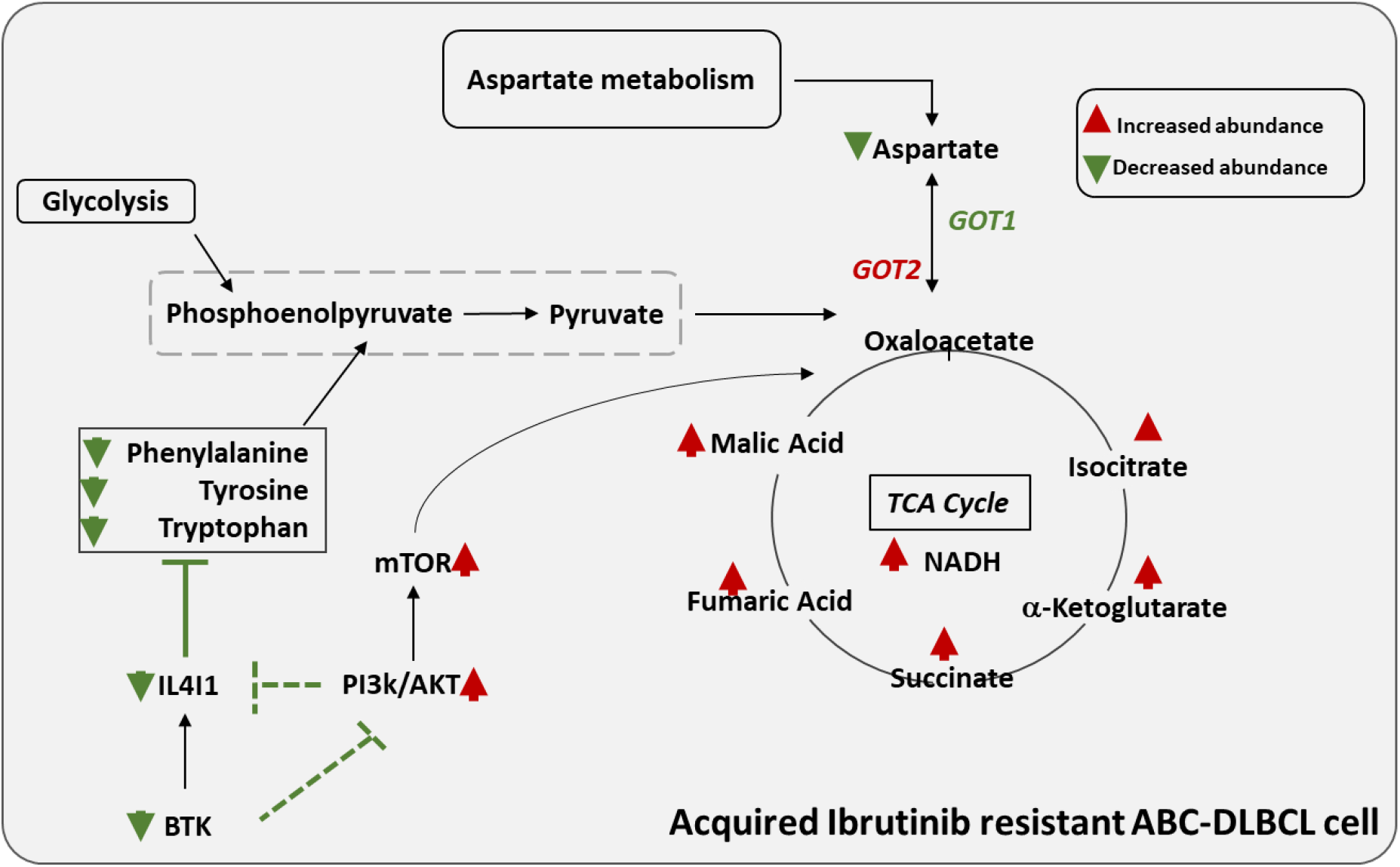
Proposed crosstalk of amino acid metabolism and TCA cycles within the Ibrutinib resistance mechanism regulated by IL4I1. The proposed metabolic crosstalks to in relation to TCA anaplerosis in drug resistant lymphoma. Modulation of BTK and PI3K AKT Pathway regulates the expression of IL4I1. Decreased IL4I1 expression affects the Phenylalanine, metabolism accompanied by decreased pyruvate levels shifting to increased oxidative phosphorylation driving robust cell proliferation in drug resistant lymphoma. One cannot exclude the possibility of ancillary metabolic pathways such as urea cycle as well as glutamate and glutamine metabolism can directly contribute to TCA intermediates fumaric acid and α-ketoglutarate, respectively.

We then explored the connection between α-ketoglutarate from the TCA cycle and the urea cycle. The intermediate metabolite can be oxidated in the presence of glutamine dehydrogenase (GDH) to produce glutamine. Glutamine and glutamate contribute to various biological processes, including nucleotide metabolism (mainly purine and pyrimidine). Also, glutamine and glutamate metabolism can feed into the arginine and proline pathway, generating more arginine to participate in the urea cycle. Urea cycle was explored on the bottom left of the figure for its ability to recycle intermediates and provide substrates for the TCA cycle. Argininosuccinate, a top VIP metabolite driving metabolic distinctiveness of the groups, plays a leading role in linking these two cycles. Argininosuccinate can replenish the TCA cycle by generating fumarate via the lyase ASL, which is mapped on the bottom left side. As argininosuccinate abundance was decreased (p-value = 0.013), we hypothesize that it will replenish the TCA cycle by generating fumarate via the lyase ASL, which is mapped on the bottom left side. Fumarate levels were increasingly abundant in addition to the upregulation of ASL. The urea cycle was perpetuated, as evidenced by the increased levels of uric acid. This finding would be expected, as dysregulated amino acid metabolism may offset the nitrogen balance in the cell.

Alternatively, there appeared to be increased activity regarding cystathionine (p-value= 0.00076), one of the top VIP metabolites driving metabolic differences between the two phenotypes. Inverse dysregulation of CBS and CTH suggest that increased cystathionine synthesis may be used for methionine salvage. Serine needed for synthesis was also more abundant in resistant cells (p-value = 0.036). This observation suggests there is an increase in methionine salvage to support cysteine levels.

## Discussion

This is the first report to perform multi-omic integration of the drug-resistant lymphoma lines reported to favor OxPhos instead of glycolysis for energy production. Many types of cancer cells exhibit pronounced metabolic reprogramming compared to non-transformed cells, as it is critical to oncogenesis. Oncogenic mutations at the core of carcinogenesis drive large-scale metabolic alterations, which are both native to cancers as well as obligatory for the transition to malignancy. The most noTable of these metabolic alterations in the activation of aerobic glycolysis, termed the Warburg effect.[29, 30] Other metabolic modifications that drive increased malignancy include activation of fatty acid biosynthesis and glutamine consumption.[31] Lymphoma metabolism has been extensively studied to identify metabolic perturbations across lymphoma subtypes.[32] DLBCL cells have been shown to metabolize more pyruvic acid, which is indicative of a glycolytic state. This finding was corroborated when levels of the glycolytic enzymes hexokinase 1 (HXK1) and phosphoglycerate kinase 1 (PGK1) were both determined to be upregulated in this lymphoma subset. Pathway analysis revealed increased expression of genes involved in glycine, serine, threonine, and pyrimidine metabolism in DLBCL. This is in line with the identification of unique metabolic profiles of lymphoma patient plasma samples from each cancer subtype.[33] Compared to other lymphomas, DLBCL patients were found to have increased circulating levels of glutamic acid and glycine, coupled with decreased levels of aspartic acid, tryptophan, and uric. These reports are not fully mirrored in our data, suggesting a unique metabolic profile of the resistant cells.

By integrating the metabolomics data with transcriptomic data, we uncovered an amino acid oxidase, IL4I1, which mediates TCA anaplerosis in Ibrutinib-resistant DLBCL (Figure 7). Research has mainly highlighted the association of IL4I1 with the regulation of immune functions, with a reliance on amino acid depletion and the formation of H2O2 and toxic phenyl-pyruvic acid.[34] BCR-induced signaling is limited by IL4I1, consequently limiting B-cell proliferation. In mouse models, in vivo experiments reveal enhanced egress of naive B cells from the bone marrow in mice with IL4I1 knockout.[35] The IL4I1 protein participates in the metabolisms of numerous amino acids including tryptophan, tyrosine, leucine, and valine.[36] However, not all of these metabolisms were significantly dysregulated in the microarray data. By connecting the alanine, aspartate, and glutamate metabolism to that of cysteine and methionine, IL4I1 forms a direct link for these two metabolic pathways to communicate. It is known that aminotransferases can facilitate amino acid-mediated TCA anaplerosis; therefore, they are considered emerging determinants of oncogenesis.[31] GOT1 and GOT2 are the respective cytoplasmic and mitochondrial varieties of glutamic-oxaloacetic transaminase enzymes (Figure 7). They participate similarly to aspartate aminotransferases (AST) by catalyzing the reversible transamination of oxaloacetate and glutamate, producing aspartate and α-ketoglutarate.[21] The mitochondrial form of glutamic-oxaloacetic transaminase, GOT2, is a prominent player in the urea cycle and TCA cycle, particularly in the malate-aspartate shuttle.[37]. The drug-resistant lymphoma lines reported in this study have been shown to favor OxPhos instead of glycolysis for energy production, so a reduction in IL4I1 expression may hold the key to this metabolic adaptation. GOT1 was found to be significantly downregulated in the HBL1 resistant cells (p-value 0.013). In TMD8R cells, inner membrane mitochondrial GOT2 was significantly downregulated (p-value 0.02), suggesting decreased transamination of aspartate to form TCA intermediate oxaloacetate (Figure 7 and supplemental Figure 5). Thus, these drug-resistant lymphomas need to recycle amino acid carbons to maintain energy production. Interestingly, studies have shown that inhibition of the electron transport chain in cells lacking GOT1 reduces cell growth, thus providing evidence for the reliance of these cancers on OxPhos metabolism.[38, 39] As such, IL4I1 cannot thus serve its function as an L-amino acid oxidase, allowing excess phenylalanine to be converted to tyrosine. Tyrosine serves as an intermediate between phenylpyruvate and phenylalanine and its reaction produces glutamate as a byproduct. With this, the cell has numerous recycling reactions that can generate glutamate for glutaminolysis. Glutamine and oxaloacetate can be respectively converted to α-ketoglutarate and aspartic acid via aspartate aminotransferase, which provide the cell a way to modulate the flux of pathways as needed.[40] It is believed that this provides a link between glycolysis, glutaminolysis, and TCA anaplerosis.[31] Conversion of glutamic acid to glutamine has been shown to fuel de novo purine biosynthesis (Figure 7 and supplemental Figure 5).[41] As such, pyrimidine metabolism and purine metabolism were among the dysregulated metabolic pathways indicating a direct effect on DNA and RNA synthesis. Our findings suggest that loss of IL4I1 induces a metabolic shift towards oxidative phosphorylation, a shared feature between the two DLBCL drug-resistant cell lines.

Our previous published study [15] suggest that loss of BTK can activate the PI3K AKT pathways; here we show that activation of PI3K AKT Pathway downregulate IL4I1 responsible for the metabolic reprogramming and drug resistance. We mapped out links that connect gene expression changes and metabolite changes to the Ibrutinib-resistant phenotype. Whether a negative feedback loop exist between the AKT and IL4I1 exist is not clearly understood, however targeting this axis using a specific inhibitor of oxidative phosphorylation or PI3K-AKT Pathway may hold the key to reverse and target the drug-induced metabolic adaptation.

## Conclusions

To conclude, our research is the first to perform multi-omic integration of the drug-resistant lymphoma lines reported to favor OxPhos instead of glycolysis for energy production via the BTK-PI3K-AKT-IL4I1 axis. Introduction of BTK restored the levels of IL4I1 and loss of IL4I1 was critical for the drug resistance, further treatment with dual inhibitor of mTOR and AKT restored IL4I1 levels, suggesting that IL4I1 is partly regulated by PI3K-AKT signaling. Aberrant expression of these genes is accompanied by a shift towards oxidative phosphorylation in these cancers, which justifies the observed TCA anaplerosis and provides a mechanism of increased metabolic activity for survival. Targeting this axis using a specific inhibitor of oxidative phosphorylation or PI3K-AKT Pathway may hold the key to reverse and target the drug-induced metabolic adaptation.

## Materials and Methods

### Cell culture/Ibrutinib resistance

DLBCL cell lines HBL1 and TMD8 were cultured in RPMI-1640 media supplemented with 10% fetal bovine serum as described previously.[15] Cultures were routinely tested for mycoplasma and cell line identities were confirmed by Short Tandem Repeat (STR) testing. The BTK inhibitor Ibrutinib (PCI-32765) was purchased from Selleckchem. Ibrutinib-resistant cells were generated as described previously. Briefly, wild-type (WT) cell lines were perpetually cultured with incremental doses of Ibrutinib for 8-10 months. Cells were passaged routinely upon confluency. However, cells passaged to less than 20 passages were utilized for experiments, after a week of drug-free culture. When cultured in the absence of Ibrutinib, all cell lines generated in this study exhibited steady Ibrutinib resistance. The wild-type BTK construct was cloned in the base vector (Lenti-X Expression System Version EF1-a; Takara Bio). STable over-expression of BTK in ibrutinib-resistant DLBCL cells was performed by using lentiviruses expressing human wild-type BTK as described previously.[15]

### IC50

To quantify surviving and/or proliferating cells, a WST-1 cell proliferation assay kit (MK400, Takara Bio) was used according to the manufacturer’s instructions. In each experiment, 10000 cells were seeded into each well of a 96-well culture dish and Ibrutinib was added the following day. Average relative absorption (OD450-OD690) was measured 72 h after the addition of Ibrutinib to estimate the number of metabolically active cells. The IC50 value was calculated using GraphPad Prism 8 software.

### Gene expression studies

Ibrutinib-resistant DLBCL cell line HBL1-R and parental line HBL1 were used in triplicate for the gene expression studies. Briefly, total RNA from each sample was quantified using the NanoDrop ND-1000 spectrophotometer. RNA integrity was assessed by standard denaturing agarose gel electrophoresis. For microarray analyses, total RNA from each sample was amplified and transcribed into fluorescent complementary RNA using the manufacturer’s Quick Amp Gene Expression Labeling Protocol, Version 5.7.9 (Agilent Technologies). The labeled complementary RNAs were hybridized onto a whole human genome oligo microarray (4 3 44K; Agilent Technologies). After washing the slides, the arrays were scanned using the Agilent microarray scanner G2505C. Agilent’s Feature Extraction software (version 11.0.1.1) was used to analyze array images. Quantile normalization and subsequent data processing were performed using the GeneSpring GX v12.1 software (Agilent Technologies). After quantile normalization of the raw data, genes that had flags in at least 3 of 24 samples were chosen for further data analyses. Differentially expressed genes were identified through fold-change and volcano filtering. Agilent’s pathway and Gene Ontology analyses were applied to determine the roles that these differentially expressed genes play in these biological pathways or Gene Ontology terms. Finally, a Venn diagram was generated to show the distinguishable gene expression profiles among samples. We deposited the corresponding raw data into the Gene Expression Omnibus data repository under accession number GSE138126.

### Immunoblotting

For immunoblotting, cells were harvested and lysed in ice-cold radio-immunoprecipitation lysis buffer (Cell Signaling Technology, Danvers, MA) containing a protease inhibitor cocktail (Roche). Equal amounts of proteins were resolved using sodium dodecyl sulfate-polyacrylamide gel electrophoresis, transferred to PVDF membrane (Bio-Rad, Hercules, CA), and analyzed with the following specific primary antibodies: anti-BTK (D3H5), anti-IL4I1 (ab-222101, Abcam), and horseradish peroxidase-conjugated anti-β-actin (A3854; Millipore Sigma, St. Louis, MO).

### Quantitative qPCR

Total RNA was isolated with the Qiagen RNeasy Plus Mini Kit following the manufacturer’s instructions. After DNase treatment, a cDNA was synthesized using a SuperScript™ III First-Strand Synthesis System (Invitrogen) using 1 μg of total RNA. Quantitative-PCR was carried out on a CFX96 Real-Time PCR System (Bio-Rad) using SYBR green PCR master mix (Applied Biosystems). Total RNA from each sample was normalized to β-actin followed by calculation of average relative quantity (RQ). The following primers were used: IL4I1 (forward primer: CGCCCGAAGACATCTACCAG, reverse primer: GATATTCCAAGAGCGTGTGCC), GOT1 (forward primer: AGTCTTTGCCGAGGTTCCG, reverse primer: GTGCGATATGCTCCCACTC), GOT2 (forward primer: GACCAAATTGGCATGTTCTGT, reverse primer: CGGCCATCTTTTGTCATGTA), ALDH7A1 (forward primer: CAACGAGCCAATAGCAAGAG, reverse primer: GCATCGCCAATCTGTCTTAC), LAP3 (forward primer: TCGGCAAAGCTCTATGGAAGT, reverse primer: GCGTCATCTCATTGGCTGG) and ALDH9A1 (forward primer: TGGAGTCAAAAATCTGGCATGG, reverse primer: AGTAGCAATTTCATCCTCCCGT).

### Metabolite extraction

Intracellular metabolites from each biological replicate were extracted using cold methanol-based extraction. Cells were harvested and counted and 1 × 106 cells were aliquoted in triplicates for metabolite extraction. Cells were washed using cold PBS before the addition of 250μL of methanol (LCMS-grade). Internal standards containing 13C and 15N labeled amino acids mix (1.2mg/mL methanol) were added to the samples at a volume of 50μL. Cells were homogenized for 2 min before incubating at −20°C for 20 min for the extraction of polar metabolites from the cells. Cells were pelleted and 150μL of supernatant was transferred to an LC-MS vial. A pooled quality control (QC) sample was collected by aliquoting an equal volume of all supernatants into a distinct vial and homogenized by vortexing. Sample vials were placed in the autosampler tray at 4°C.

### LC-MS/MS system

The LC-MS/MS analyses were performed on a Vanquish ultra-high-performance liquid chromatography (UHPLC) system (Thermo Scientific, Waltham MA, USA) coupled to a QExactive™ Hybrid Quadrupole-Orbitrap™ Mass Spectrometer (Thermo Scientific, Waltham MA, USA). A sample volume of 5 μL was injected onto an XBridge BEH Amide XP Column, 130Å (150 mm × 2.1 mm ID, particle size 2.5 μm) (Waters Corporation, Milford, MA, USA). The column oven was maintained at 40°C. Mobile phase A consisted of a mixture of 5 mM NH4Ac in ACN/H2O (10:90, v/v) with 0.1% acetic acid. Mobile phase B consisted of 5 mM NH4Ac in ACN/H2O (90:10, v/v) with 0.1% acetic acid. The mobile phases were delivered at a flow rate of 0.3 mL/min for a 20 min run with the following stepwise gradient for solvent B: firstly 70%; 0-5 min 30%; 5-9 min 30%; 9-11 min 70%. A divert valve was used to direct the flow to waste during the final 5 min of the run. The QExactive™ was equipped with an electrospray ionization source (ESI) that was operated in both negative and positive ion modes to encompass a broad range of metabolite detection. The (ESI) source setting and the compound dependent scan conditions were optimized for full scan MS mode and ranged between 150 and 2000m/z. The ion spray voltage was set at 4kV with a capillary temperature of 320°C. The sheath gas rate was set to 10 arbitrary units. Scans of 1 ms were performed at 35,000 units resolution.

MS-based untargeted metabolomics was conducted in triplicates to reveal the metabolic modifications driving increased cell survival and robust proliferation in the drug-resistant phenotype. The analysis included pooled quality control samples at the beginning and end of the run, in addition to a QC followed by a blank among every 10 biological sample injections. The pooled samples were also utilized for the top 10 MS/MS analysis with dynamic exclusion during the analysis for compound identification in the analysis.

### Data processing and statistical analysis

All raw UHPLC−MS data were imported into the automatic feature annotation and interpretation software Compound Discoverer 3.1 (Thermo Scientific, Waltham MA, USA) for metabolite identification. The MS data were searched against our in-house database containing experimentally obtained MS/MS spectra of 171 authentic analytical standards, and several online databases including KEGG (http://www.kegg.com/), the human metabolite database (http://www.hmdb.ca/), ChemSpider (http://www.chemspider.com/) and PubChem compound database (http://ncbi.nim.nih.gov/) for metabolite identification. The collected data were then normalized to cell count per replicate and spectra were filtered to reduce redundancy and to ensure instrument reproducibility. Any metabolite with a coefficient of variance greater than 30% was removed before further analysis. Statistical analyses, including univariate (T-test) and multivariate (PLS-DA) and pathway analysis, were conducted using the online resource MetaboAnalyst 4.0.[42] The global test packages examine metabolites within respective pathways to determine their association with an experimental variable. Top pathways were selected based on pathway impact > 0.2 and – log (p) > 2. Primary metabolites from top altered pathways were selected for further analysis. PLS-DA was used for the interpretation of the metabolic differences between the sensitive and resistant cell lines. VIP plots were generated to observe top metabolites contributing to metabolic uniqueness.

### Gene-Metabolite interaction map

Gene-Metabolite interaction networks are created within MetaboAnalyst by mapping the annotated metabolites or genes to a comprehensive gene-metabolite interaction data on STITCH (‘search tool for interactions of chemicals’).[43] Using interaction data from published peer-reviewed literature, a search algorithm can identify metabolites that immediately interact with another given compound or gene and subsequently map the nodes as direct neighbors in the network. MetaboAnalyst assesses degree centrality and betweenness centrality to determine node importance within the network. In the interaction map, the number of connections between a node and others determines its degree of centrality. The betweenness centrality calculates the number of shortest routes going through the node. These measures allow for the identification of nodes acting as hubs in the network.

## Supplementary Materials

The gene expression dataset is deposited to the Gene Expression Omnibus data repository under accession number GSE138126. The Mass spectrometry raw data are available on request analysed data is present in supplemental files. All the cell lines used and generated will be available upon request.

## Author Contributions

Study concept design: JZ, LS, and FC. Acquisition, analysis, or interpretation of data: all authors. Statistical analysis: FC. Critical revision of the manuscript for important intellectual content: all authors. All authors read and approved the final manuscript.

## Acknowledgments

We thank Dr Jennifer Woyach for kindly providing the wild type BTK constructs.

## Bibliography

1. Armitage, J.O., My treatment approach to patients with diffuse large B-cell lymphoma. Mayo Clin Proc, 2012. 87(2): p. 161–71.

2. Cultrera, J.L. and S.M. Dalia, Diffuse large B-cell lymphoma: current strategies and future directions. Cancer Control, 2012. 19(3): p. 204–13.

3. Roschewski, M., L.M. Staudt, and W.H. Wilson, Diffuse large B-cell lymphoma-treatment approaches in the molecular era. Nat Rev Clin Oncol, 2014. 11(1): p. 12–23.

4. Lenz, G., et al., Stromal gene signatures in large-B-cell lymphomas. N Engl J Med, 2008. 359(22): p. 2313–23.

5. Lenz, G., Insights into the Molecular Pathogenesis of Activated B-Cell-like Diffuse Large B-Cell Lymphoma and Its Therapeutic Implications. Cancers (Basel), 2015. 7(2): p. 811–22.

6. Davis, R.E., et al., Chronic active B-cell-receptor signalling in diffuse large B-cell lymphoma. Nature, 2010. 463(7277): p. 88–92.

7. Boukhiar, M.A., et al., Targeting early B-cell receptor signaling induces apoptosis in leukemic mantle cell lymphoma. Exp Hematol Oncol, 2013. 2(1): p. 4.

8. Tucker, D.L. and S.A. Rule, A critical appraisal of ibrutinib in the treatment of mantle cell lymphoma and chronic lymphocytic leukemia. Ther Clin Risk Manag, 2015. 11: p. 979–90.

9. Jones, J., et al., Evaluation of 230 patients with relapsed/refractory deletion 17p chronic lymphocytic leukaemia treated with ibrutinib from 3 clinical trials. Br J Haematol, 2018. 182(4): p. 504–512.

10. Wilson, W.H., et al., Targeting B cell receptor signaling with ibrutinib in diffuse large B cell lymphoma. Nat Med, 2015. 21(8): p. 922–6.

11. Rudelius, M., et al., Inhibition of focal adhesion kinase overcomes resistance of mantle cell lymphoma to ibrutinib in the bone marrow microenvironment. Haematologica, 2018. 103(1): p. 116–125.

12. Pera, B., et al., Metabolomic Profiling Reveals Cellular Reprogramming of B-Cell Lymphoma by a Lysine Deacetylase Inhibitor through the Choline Pathway. EBioMedicine, 2018. 28: p. 80–89.

13. Caro, P., et al., Metabolic signatures uncover distinct targets in molecular subsets of diffuse large B cell lymphoma. Cancer Cell, 2012. 22(4): p. 547–60.

14. Xiong, J., et al., MYC is a positive regulator of choline metabolism and impedes mitophagy-dependent necroptosis in diffuse large B-cell lymphoma. Blood Cancer J, 2017. 7(7): p. e0.

15. Jain, N., et al., Targeting phosphatidylinositol 3 kinase-beta and -delta for Bruton tyrosine kinase resistance in diffuse large B-cell lymphoma. Blood Adv, 2020. 4(18): p. 4382–4392.

16. Kuo, H.-P., et al., Combination of Ibrutinib and ABT-199 in Diffuse Large B-Cell Lymphoma and Follicular Lymphoma. Molecular Cancer Therapeutics, 2017. 16(7): p. 1246–1256.

17. Pal Singh, S., F. Dammeijer, and R.W. Hendriks, Role of Bruton’s tyrosine kinase in B cells and malignancies. Molecular Cancer, 2018. 17(1).

18. Phan, L.M., S.C. Yeung, and M.H. Lee, Cancer metabolic reprogramming: importance, main features, and potentials for precise targeted anti-cancer therapies. Cancer Biol Med, 2014. 11(1): p. 1–19.

19. Smilde, A.K., et al., Dynamic metabolomic data analysis: a tutorial review. Metabolomics, 2010. 6(1): p. 3–17.

20. Boulland, M.-L., et al., Human IL4I1 is a secreted l-phenylalanine oxidase expressed by mature dendritic cells that inhibits T-lymphocyte proliferation. Blood, 2007. 110(1): p. 220–227.

21. Jiang, X., et al., Recombinant expression, purification and crystallographic studies of the mature form of human mitochondrial aspartate aminotransferase. Biosci Trends, 2016. 10(1): p. 79–84.

22. Kozlov, A.M., et al., Lactate preconditioning promotes a HIF-1alpha-mediated metabolic shift from OXPHOS to glycolysis in normal human diploid fibroblasts. Sci Rep, 2020. 10(1): p. 8388.

23. Hanahan, D. and R.A. Weinberg, Hallmarks of cancer: the next generation. Cell, 2011. 144(5): p. 646–74.

24. Khapare, N., et al., Plakophilin3 loss leads to an increase in PRL3 levels promoting K8 dephosphorylation, which is required for transformation and metastasis. PLoS One, 2012. 7(6): p. e38561.

25. Zijun Y. Xu-Monette, S.Z., Xin Li1, Ganiraju C. Manyam, Xiao-xiao Wang, Yi Xia, Carlo Visco, Alexandar Tzankov Miguel A. Piris6, L. Jeffrey Medeiros1, and Ken H. Young, p63 expression confers significantly better survival outcomes in high-risk diffuse large B-cell lymphoma and demonstrates p53-like and p53-independent tumor suppressor function. AGING, 2016. 8(2).

26. Qing Ye, Z.Y. X.-M., Alexandar Tzankov, Lijuan Deng, Xiaoxiao Wang, Ganiraju C. Manyam, Carlo Visco, Eric D. Hsi, Michael B. Møller, Miguel A. Piris, Jane N. Winter, L. Jeffrey Medeiros1 Shimin Hu1 and Ken H. Young, Prognostic impact of concurrent MYC and BCL6 rearrangements and expression in de novo diffuse large B-cell lymphoma. Oncotarget, 2015. 7.

27. Siddiqui, A. and P. Ceppi, A non-proliferative role of pyrimidine metabolism in cancer. Mol Metab, 2020. 35: p. 100962.

28. Molina, J.R., et al., An inhibitor of oxidative phosphorylation exploits cancer vulnerability. Nat Med, 2018. 24(7): p. 1036–1046.

29. Warburg, O., On the Origin of Cancer Cells. Science, 1956. 123(3191): p. 309–314.

30. Liberti, M.V. and J.W. Locasale, The Warburg Effect: How Does it Benefit Cancer Cells? Trends Biochem Sci, 2016. 41(3): p. 211–218.

31. Smith, B., et al., Addiction to Coupling of the Warburg Effect with Glutamine Catabolism in Cancer Cells. Cell Reports, 2016. 17(3): p. 821–836.

32. Schwarzfischer, P., et al., Comprehensive Metaboproteomics of Burkitt’s and Diffuse Large B-Cell Lymphoma Cell Lines and Primary Tumor Tissues Reveals Distinct Differences in Pyruvate Content and Metabolism. J Proteome Res, 2017. 16(3): p. 1105–1120.

33. Barberini, L., et al., The Metabolomic Profile of Lymphoma Subtypes: A Pilot Study. Molecules, 2019. 24(13).

34. Molinier-Frenkel, V., A. Prevost-Blondel, and F. Castellano, The IL4I1 Enzyme: A New Player in the Immunosuppressive Tumor Microenvironment. Cells, 2019. 8(7).

35. Bod, L., et al., IL-4-Induced Gene 1: A Negative Immune Checkpoint Controlling B Cell Differentiation and Activation. J Immunol, 2018. 200(3): p. 1027–1038.

36. Serrano-Carbajal, E.A., J. Espinal-Enriquez, and E. Hernandez-Lemus, Targeting Metabolic Deregulation Landscapes in Breast Cancer Subtypes. Front Oncol, 2020. 10: p. 97.

37. Yang, H., et al., SIRT3-dependent GOT2 acetylation status affects the malate-aspartate NADH shuttle activity and pancreatic tumor growth. EMBO J, 2015. 34(8): p. 1110–25.

38. Birsoy, K., et al., An Essential Role of the Mitochondrial Electron Transport Chain in Cell Proliferation Is to Enable Aspartate Synthesis. Cell, 2015. 162(3): p. 540–551.

39. Deberardinis, R.J., A mitochondrial power play in lymphoma. Cancer Cell, 2012. 22(4): p. 423–4.

40. Altman, B.J., Z.E. Stine, and C.V. Dang, From Krebs to clinic: glutamine metabolism to cancer therapy. Nat Rev Cancer, 2016. 16(10): p. 619–34.

41. Tardito, S., et al., Glutamine synthetase activity fuels nucleotide biosynthesis and supports growth of glutamine-restricted glioblastoma. Nature Cell Biology, 2015. 17(12): p. 1556–1568.

42. Chong, J., D.S. Wishart, and J. Xia, Using MetaboAnalyst 4.0 for Comprehensive and Integrative Metabolomics Data Analysis. Curr Protoc Bioinformatics, 2019. 68(1): p. e86.

43. Szklarczyk, D., et al., STITCH 5: augmenting protein-chemical interaction networks with tissue and affinity data. Nucleic Acids Res, 2016. 44(D1): p. D380–4.

