## Supplementary material for "Multiomics Integration Elucidates Metabolic Modulators of Drug Resistance in Lymphoma"

**Multiomics Integration Elucidates Onco-Metabolic Modulators of Drug Resistance in Activated B-Cell Lymphoma**

### Supplemental data

A

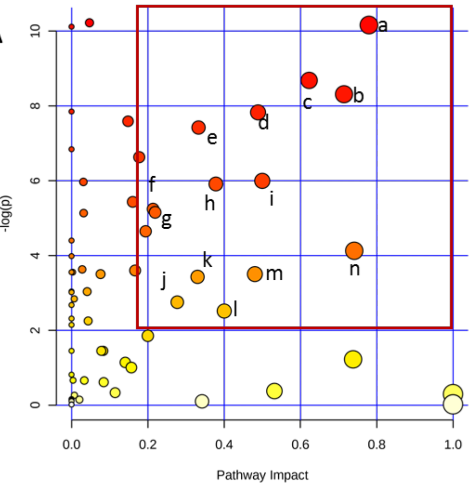


| | |
|---|---|
| a | Alanine, aspartate and glutamate metabolism |
| b | Taurine and hypotaurine metabolism |
| c | Histidine metabolism |
| d | Glycine, serine and threonine metabolism |
| e | Pyrimidine metabolism |
| f | Purine metabolism |
| g | Glutathione metabolism |
| h | Cysteine and methionine metabolism |
| i | Glutamine and glutamate metabolism |
| j | Glycerophospholipid metabolism |
| k | Arginine and proline metabolism |
| l | Tyrosine Metabolism |
| m | Starch and sucrose metabolism |
| n | Arginine biosynthesis |

B

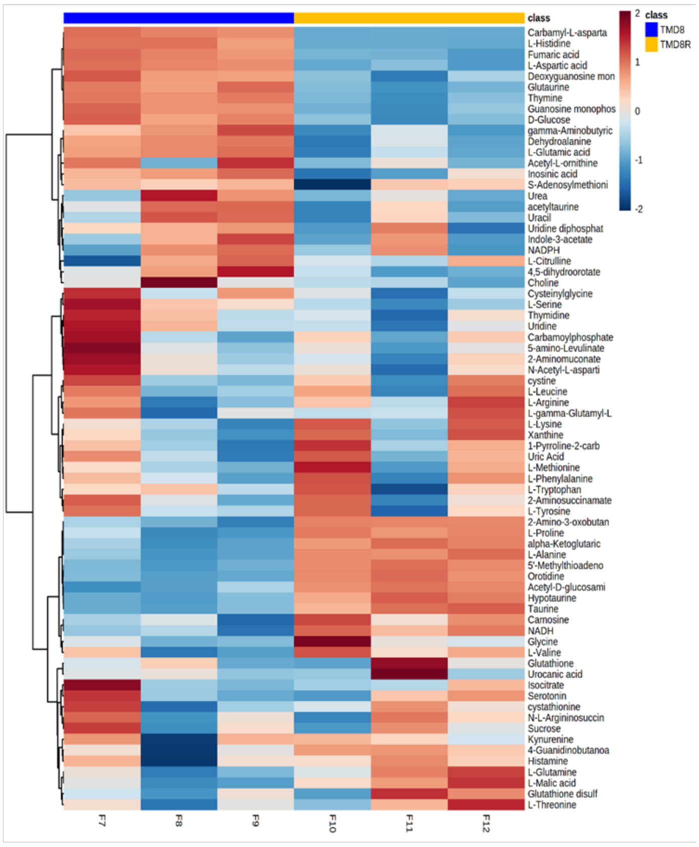


**Supplemental Figure 1:** Validation of similar metabolic profile in TMD8 A) Summary of main altered pathways based on untargeted metabolomics analysis of each cell type. B) Heatmap of the seventy-four metabolites detected from dysregulated pathways

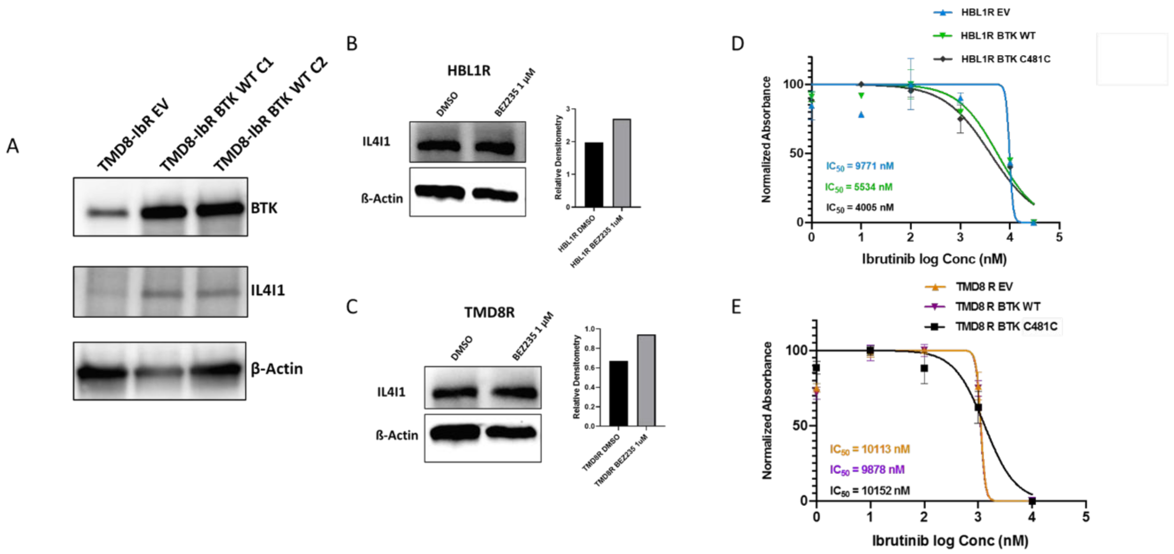


**Supplemental figure 2:** BTK Rescue experiments altered IL4I1 expression A) IL4I1 western blot showing expression across diverse lymphoma mutants. Total IL4I1 and BTK protein levels in BTK rescued B) HBL1R and C) TMD8R upon treatment with PI3K selective as well as PI3K/mTOR inhibitors. D-E) IC50 graphs among resistant lymphomas exhibiting variable BTK expression.

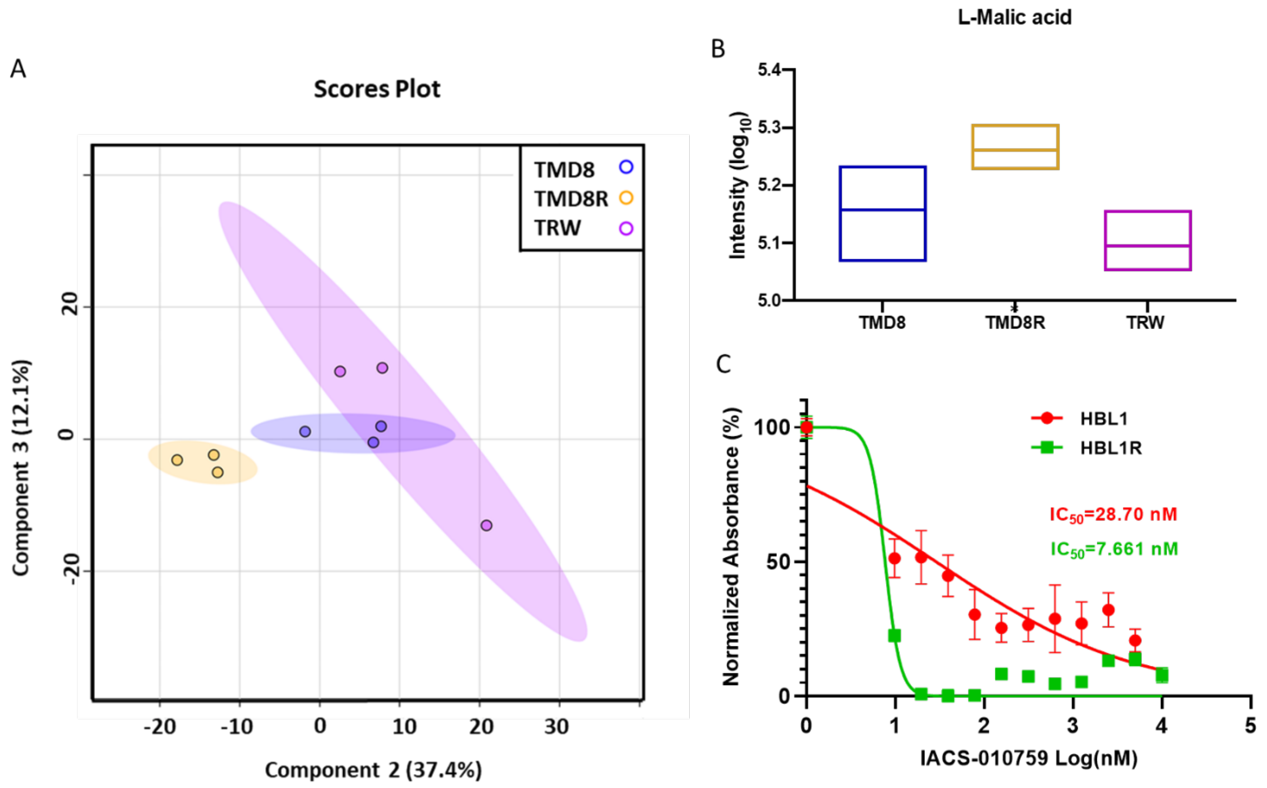


**Supplemental figure 3:** Partial reversal trend in TMD8R cells upon overexpression of WT BTK gene A) PLS-DA plot B) Relative abundance change of TCA metabolites in TMD8, TMD8R and TMD8R expressing WT BTK gene. C) IC-50 of the IACS-010759 showing more sensitivity in the ibrutinib resistant HBL1R.

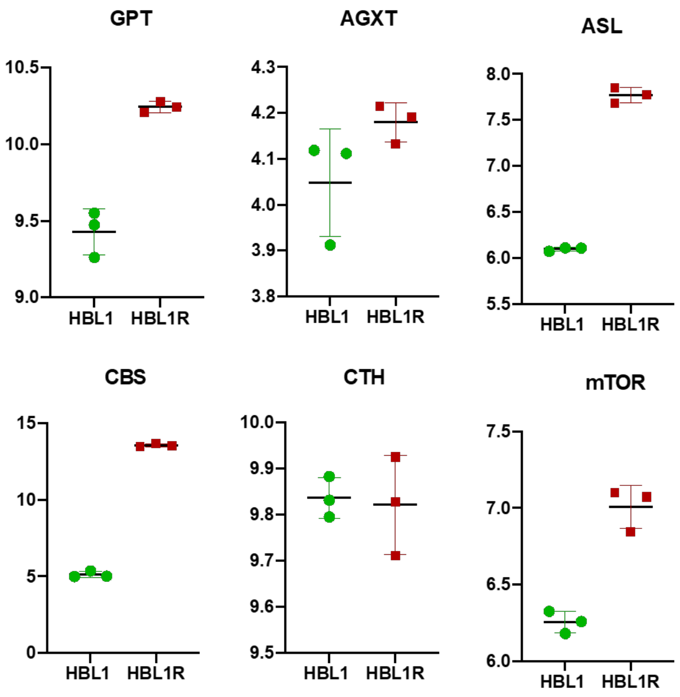


***Supplemental figure 4****: Fold change expression of genes in the proposed crosstalk*

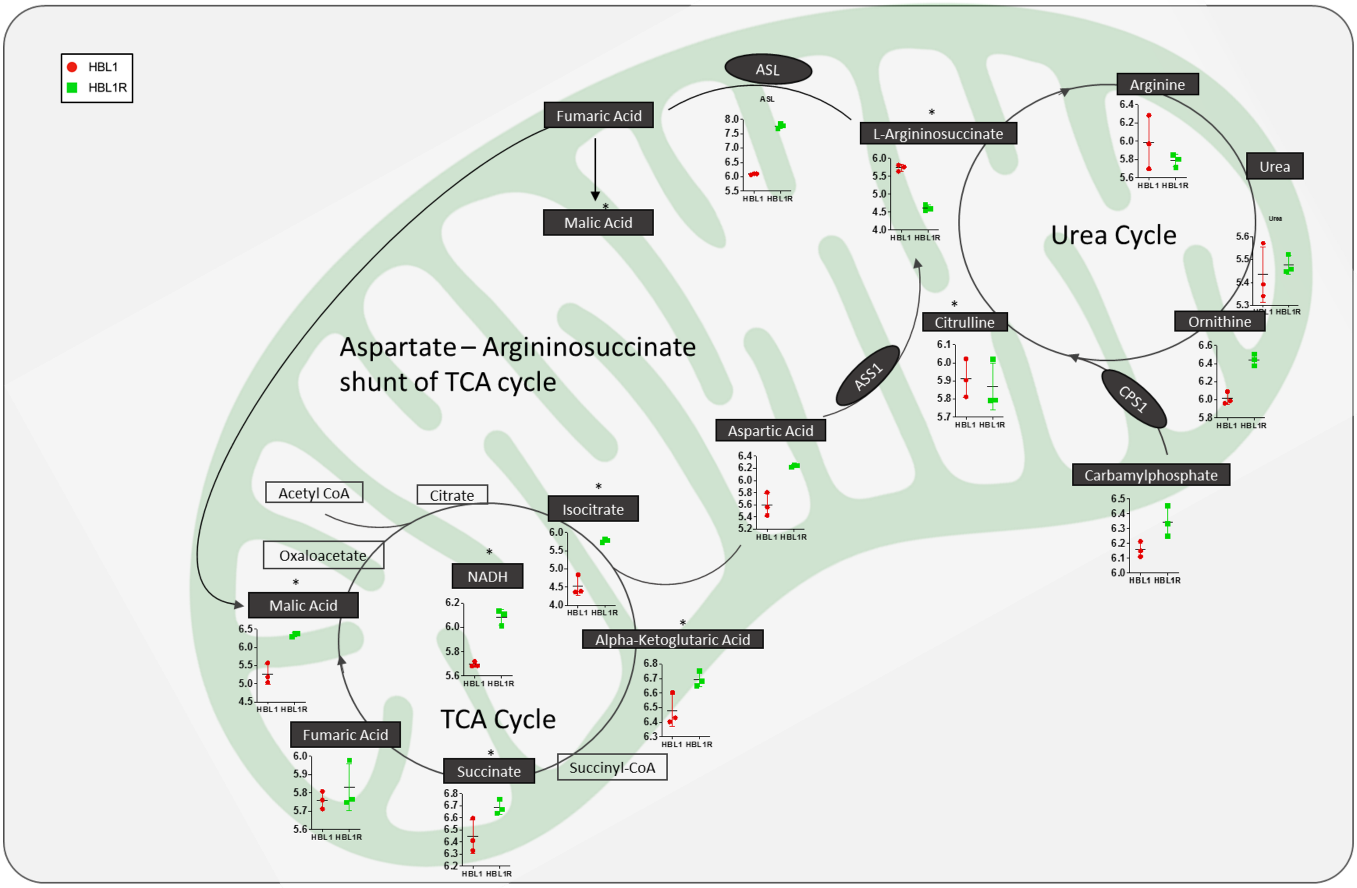


**Supplemental figure 5:** Aspartate-arginino-succinate shunt at the crosstalk of the TCA and urea cycles constructed based on metabolomics analysis of the HBL1/R cell pair.

**Supplemental Tables**

**Supplemental Table 1:** Metabolites of the significantly altered metabolic pathways in the HBL1R phenotype.

| Name | Abundance Ratio |
|---|---|
| 1-Pyrroline-2-carboxylate | 1.08 |
| 2-Amino-3-oxobutanoic acid | 1.09 |
| 2-Aminomuconate | 1.09 |
| 2-Aminosuccinamate | 1.09 |
| 4-Aminobutanoate | 0.97 |
| 4-Guanidinobutanoate | 0.97 |
| 4,5-dihydroorotate | 1.09 |
| 5-amino-Levulinate | 1.10 |
| 5'-Methylthioadenosine | 1.42 |
| Acetyl-L-ornithine | 0.97 |
| Acetyltaurine | 1.00 |
| alpha-Ketoglutaric acid | 1.03 |
| Carbamoylphosphate | 1.03 |
| Carbamyl-L-aspartate | 1.09 |
| Carnosine | 1.00 |
| Choline | 0.96 |
| Cystathionine | 1.42 |
| Cysteinylglycine | 1.09 |
| Cystine | 0.96 |
| Dehydroalanine | 0.98 |

| | |
|---|---|
| Deoxyguanosine monophosphate | 0.96 |
| Fumaric acid | 1.01 |
| Glutathione | 1.17 |
| Glutathione disulfide | 1.20 |
| Glutaurine | 0.90 |
| Glycine | 1.01 |
| Guanosine monophosphate | 0.90 |
| Histamine | 0.98 |
| Hypotaurine | 1.08 |
| Indole-3-acetate | 1.05 |
| Inosinic acid | 0.97 |
| Isocitrate | 1.28 |
| Kynurenine | 0.98 |
| L-Alanine | 1.04 |
| L-Arginine | 0.97 |
| L-Aspartic acid | 1.11 |
| L-Citrulline | 0.99 |
| L-gamma-Glutamyl-L-leucine | 1.01 |
| L-Glutamic acid | 0.99 |
| L-Glutamine | 0.97 |
| L-Histidine | 1.01 |
| L-Leucine | 0.97 |
| L-Lysine | 1.01 |
| L-Malic acid | 1.21 |
| L-Methionine | 1.00 |

| | |
|---|---|
| L-Phenylalanine | 0.99 |
| L-Proline | 0.98 |
| L-Serine | 1.03 |
| L-Threonine | 0.88 |
| L-Tryptophan | 0.99 |
| L-Tyrosine | 0.99 |
| L-Valine | 0.98 |
| N-(L-Arginino)succinate | 0.80 |
| N-Acetyl-L-aspartic acid | 1.05 |
| NADH | 1.07 |
| NADPH | 0.90 |
| Orotidine | 1.07 |
| S-Adenosylmethionine | 1.00 |
| Serotonin | 0.97 |
| Taurine | 1.05 |
| Thymidine | 1.09 |
| Thymine | 1.12 |
| Uracil | 0.99 |
| Urea | 1.01 |
| Uric Acid | 1.03 |
| Uridine | 1.08 |
| Urocanic acid | 1.10 |
| Xanthine | 1.06 |

**Supplemental Table 2**: Altered genes from alanine, aspartate, and glutamate metabolic pathway.

| SYMBOL | RANK METRIC SCORE | RUNNING ES |
|---|---|---|
| ASL | -0.60 | -0.02 |
| ALDH5A1 | -1.39 | 0.03 |
| GPT2 | -0.21 | 0.05 |
| GOT2 | -0.26 | 0.06 |
| PPAT | -0.22 | 0.06 |
| GLS2 | -0.14 | 0.09 |
| ACY3 | -0.11 | 0.10 |
| NIT2 | -0.10 | 0.10 |
| AGXT | -0.08 | 0.11 |
| CAD | -0.05 | 0.14 |
| GLUD1 | -0.03 | 0.17 |
| GLUD2 | 0.01 | 0.21 |
| ASS1 | 3.32 | 0.27 |
| GAD1 | 0.11 | 0.36 |
| ALDH4A1 | 0.10 | 0.36 |
| CPS1 | 1.44 | 0.36 |
| ADSL | 0.15 | 0.40 |
| IL4I1 | 0.99 | 0.43 |
| GLUL | 0.85 | 0.49 |
| ABAT | 0.55 | 0.49 |
| GOT1 | 0.35 | 0.50 |
| ADSS | 0.28 | 0.51 |

| | | |
|---|---|---|
| GFPT1 | 0.47 | 0.51 |
| GLS | 0.34 | 0.53 |

**Supplemental Table 3**: Altered genes from cysteine and methionine metabolic pathway.

| SYMBOL | RANK METRIC SCORE | RUNNING ES |
|---|---|---|
| TRDMT1 | -0.83 | -0.41 |
| LDHA | -0.22 | -0.40 |
| APIP | -0.24 | -0.39 |
| ENOPH1 | -0.31 | -0.38 |
| GOT2 | -0.26 | -0.38 |
| LDHC | -0.14 | -0.37 |
| AMD1 | -0.15 | -0.36 |
| MTAP | -0.84 | -0.34 |
| DNMT3B | -1.31 | -0.27 |
| CTH | 0.00 | -0.23 |
| SRM | 0.02 | -0.19 |
| LDHB | 0.03 | -0.17 |
| DNMT3L | -1.60 | -0.15 |
| LDHAL6A | 0.09 | -0.11 |
| MTR | 0.11 | -0.09 |
| AHCY | 0.10 | -0.09 |
| MAT2B | 0.13 | -0.08 |
| SDS | 0.13 | -0.07 |
| MAT2A | 0.21 | -0.03 |

| | | |
|---|---|---|
| DNMT3A | 0.19 | -0.02 |
| AHCYL1 | 0.21 | -0.01 |
| CBS | -2.26 | 0.01 |
| IL4I1 | 0.99 | 0.03 |
| GOT1 | 0.35 | 0.05 |
| ADI1 | 0.43 | 0.05 |
| MPST | 0.74 | 0.07 |

**Supplemental Table 4**: Metabolites of the significantly altered metabolic pathways in TMD8R phenotype.

| Name | Abundance Ratio |
|---|---|
| 1-Pyrroline-2-carboxylate | 1.40 |
| 2-Amino-3-oxobutanoic acid | 2.40 |
| 2-Aminomuconate | 0.47 |
| 2-Aminosuccinamate | 0.98 |
| 4,5-dihydroorotate | 0.39 |
| 4-Guanidinobutanoate | 1.16 |
| 5-amino-Levulinate | 0.53 |
| 5'-Methylthioadenosine | 1.31 |
| Acetyl-L-ornithine | 0.91 |
| acetyltaurine | 0.40 |
| alpha-Ketoglutaric acid | 2.33 |
| Carbamoylphosphate | 0.95 |
| Carbamyl-L-aspartate | 0.18 |

| | |
|---|---|
| Carnosine | 1.71 |
| Choline | 0.74 |
| Choline | 0.74 |
| Cysteinylglycine | 0.65 |
| cystine | 0.94 |
| Dehydroalanine | 0.62 |
| Deoxyguanosine monophosphate | 0.33 |
| D-Glucose | 0.45 |
| Fumaric acid | 0.24 |
| gamma-Aminobutyric acid | 0.52 |
| Glutathione | 1.31 |
| Glutathione disulfide | 1.08 |
| Glutaurine | 0.08 |
| Glycine | 1.74 |
| Guanosine monophosphate | 0.26 |
| Histamine | 1.13 |
| Hypotaurine | 1.72 |
| Indole-3-acetate | 0.50 |
| Inosinic acid | 0.66 |
| Isocitrate | 0.96 |
| Kynurenine | 1.04 |
| L-Alanine | 1.79 |
| L-Arginine | 1.30 |
| L-Aspartic acid | 0.21 |
| L-Citrulline | 0.92 |
| L-Cystathionine | 1.08 |

| | |
|---|---|
| L-gamma-Glutamyl-L-leucine | 1.11 |
| L-Glutamic acid | 0.61 |
| L-Glutamine | 1.32 |
| L-Histidine | 0.04 |
| L-Leucine | 1.11 |
| L-Lysine | 1.51 |
| L-Malic acid | 1.66 |
| L-Methionine | 1.26 |
| L-Phenylalanine | 1.12 |
| L-Proline | 1.32 |
| L-Serine | 0.69 |
| L-Threonine | 2.00 |
| L-Tryptophan | 0.99 |
| L-Tyrosine | 0.98 |
| L-Valine | 1.68 |
| N-(L-Arginino)succinate | 0.99 |
| N4-(beta-N-Acetyl-D-glucosaminyl)-L-asparagine | 2.86 |
| N-Acetyl-L-aspartic acid | 0.44 |
| NADH | 1.41 |
| NADPH | 0.55 |
| Orotidine | 1.96 |
| S-Adenosylmethionine | 0.64 |
| Serotonin | 0.89 |
| Sucrose | 0.64 |
| Taurine | 1.34 |

| | |
|---|---|
| Thymidine | 0.30 |
| Thymine | 0.50 |
| Uracil | 0.20 |
| Urea | 0.64 |
| Uric Acid | 1.17 |
| Uridine | 0.25 |
| Uridine diphosphate glucose | 0.73 |
| Urocanic acid | 2.39 |
| Xanthine | 1.66 |
